# The UPR^MT^ is a critical mediator of nutrient immunity

**DOI:** 10.64898/2026.07.11.737880

**Authors:** Ayane Maruichi, Hanlin Zhang, Alexandra May Viret, Larry K. Joe, Andrew Dillin

## Abstract

Mitochondria serve as critical signaling hubs that monitor the intracellular environment to orchestrate cellular stress responses during pathogen infection. The mitochondrial unfolded protein response (UPR^MT^), a key surveillance pathway, is widely considered to be activated by pathogen-derived toxins that directly damage host mitochondria. In contrast to this prevailing model, we demonstrate that *E. faecalis,* which lacks specialized mitochondrial toxins, activates the UPR^MT^ in *C. elegans* via a combination of metabolic and oxidative stress. As both *C. elegans* and *E. faecalis* are heme auxotrophs, *E. faecalis*-ingested animals result in severe heme deficiency, leading to the disruption of the electron transport chain and activation of the UPR^MT^. During live infection, this primary mitochondrial stress is further amplified due to the Fenton reaction, driven by host NADPH oxidase BLI-3-generated hydrogen peroxide. Ultimately, our findings position the UPR^MT^ as a fundamental homeostatic sensor that monitors metabolic and oxidative imbalances at the host-pathogen interface.

## Introduction

Organisms must continually adapt to dynamic environmental stressors to maintain cellular and physiological homeostasis. Pathogenic bacterial infection is a particularly complex challenge, as it initiates a “host-pathogen arms race” in which pathogens attempt to hijack or deplete essential host resources, while hosts activate immune defenses. As mitochondria are central hubs of cellular metabolism and innate immunity, they are often primary targets during bacterial infection.^1,2^ Consequently, diverse bacterial pathogens have evolved sophisticated strategies to sabotage host mitochondria and enhance their own survival. For example, *Listeria monocytogenes* and *Helicobacter pylori* produce pore-forming toxins, Listeriolysin O (LLO) and vacuolating cytotoxin A (VacA), respectively, that trigger rapid mitochondrial fission and collapse of the mitochondrial membrane potential.^3,4^ Additionally, other pathogens possess electron transport chain (ETC) inhibitors such as Antimycin A of *Streptomyces*^5^ and iron chelators such as pyoverdine of *Pseudomonas aeruginosa* (*P. aeruginosa*)^6^ to directly perturb organellar proteostasis. Ultimately, these mitochondrial insults impair host cellular functions, disrupting the delicate homeostatic balance crucial for organismal survival.

To encounter these impairments, host organisms utilize surveillance mechanisms to monitor mitochondrial health and maintain cellular homeostasis. One of the key quality-control programs is the mitochondrial unfolded protein response (UPR^MT^), a transcriptional response activated upon mitochondrial stress to restore organellar function and promote organismal health. The UPR^MT^ is highly conserved from *Caenorhabditis elegans* (*C. elegans*) to mammals, and this response is primarily governed by the transcription factor ATFS-1 in *C. elegans*. Under mitochondrial stress, ATFS-1 translocates to the nucleus to upregulate genes, including mitochondrial chaperones and proteases, which help restore organellar function.^7,8,9^ Beyond its role in protein quality control, UPR^MT^ activation directly induces immune response genes and protects animals during bacterial infection.^10,11^

This UPR^MT^-mediated innate immune activation was first reported in *C. elegans* infected with bacterial pathogen *P. aeruginosa*, and it has served as the primary model for studying the mechanism of bacteria-induced UPR^MT^. Extensive characterization has revealed several bacterial toxins and metabolites of *P. aeruginosa*, such as pyocyanin and hydrogen cyanide, that directly inhibit the ETC and damage mitochondria, thereby activating the UPR^MT^.^10,12,13^ However, despite the diversity of mitochondria-targeting strategies used by various bacterial pathogens, our mechanistic understanding of UPR^MT^ activation during infection relies heavily on this *C. elegans-P. aeruginosa* model. Therefore, it remains largely unexplored whether this ‘bacterial virulence-based’ mechanism serves as a universal model for sensing diverse bacterial pathogens and activating UPR^MT^.

In this study, we report that the opportunistic pathogen *Enterococcus faecalis* (*E. faecalis*) activates the UPR^MT^ in *C. elegans* despite the absence of specialized mitochondria-targeting toxins.^14,15^ *E. faecalis* is a Gram-positive commensal of the mammalian gastrointestinal tract that frequently emerges as an opportunistic pathogen and a leading cause of nosocomial infections. Particularly in immunocompromised hosts, it causes a broad range of disease, from urinary tract infections to life-threatening endocarditis.^16,17^ Mechanistically, we reveal that this response is driven by the host-centric endogenous response to dietary heme deficiency and oxidative stress. Specifically, because *C. elegans* is a heme-auxotrophic animal that relies entirely on its bacterial diet for heme acquisition, the ingestion of *E. faecalis,* which is also a heme auxotroph, creates a severe heme-deficient state in the host. Since heme is an essential cofactor for the ETC, this heme deficiency acts as a primary mitochondrial stressor elicited by both live and dead bacteria. Additionally, during live *E. faecalis* infection, hydrogen peroxide (H_2_O_2_) generated by the host NADPH oxidase Bli-3 fuels the Fenton reaction, producing highly reactive hydroxyl radicals within the mitochondria and amplifying mitochondrial damage. Our findings provide a new perspective that mitochondrial surveillance during infection is not exclusively triggered by specialized bacterial virulence factors. Instead, we propose a model in which the host monitors the metabolic and oxidative landscape at the host-pathogen interface, positioning UPR^MT^ as a fundamental homeostatic sensor that detects cellular imbalances and maintains organismal health.

## Results

### *E. faecalis* infection induces mitochondrial dysfunction and UPR^MT^ activation in *C. elegans*

To investigate the mechanisms of UPR^MT^ activation in the context of host-pathogen interactions beyond the well-characterized *P. aeruginosa* model, we screened for novel bacterial species that impact host mitochondrial proteostasis. Using the *hsp-6p*::GFP transcriptional reporter, a mitochondrial chaperone highly induced by UPR^MT^ activation^17^, we found that *C. elegan*s larvae reared on the opportunistic pathogen *E. faecalis* OG1RF exhibit robust and systemic activation of the UPR^MT^ (Fig. 1A). Consistent with previous findings, these *E. faecalis*-grown animals also exhibited a severe developmental arrest phenotype, failing to progress beyond the L2 stage.^18^ As these arrested animals are metabolically compromised and unsuitable for further characterization of mitochondrial functions, we utilized a bacterial titration strategy. By mixing *E. faecalis* with the standard laboratory diet *E. coli* HT115 at varying ratios, we successfully decoupled the UPR^MT^ activation from the growth arrest phenotype (Fig. 1B). This approach allowed animals to reach reproductive adulthood while maintaining dose-dependent UPR^MT^ induction (Fig.1C). This *E. faecalis*-mediated UPR^MT^ was further validated by a dose-dependent increase in DVE-1 nuclear localization (Fig. 1D) and was characterized to be activated in an *atfs-1*-dependent manner (Fig. S1A). Notably, *E. faecalis* exposure did not activate the endoplasmic reticulum unfolded protein response (UPR^ER^) or the cytosolic heat stress response (HSR), suggesting the proteotoxic stress induced by *E. faecalis* is highly specific to mitochondria (Fig. S1B, S1C).

**Figure 1.**
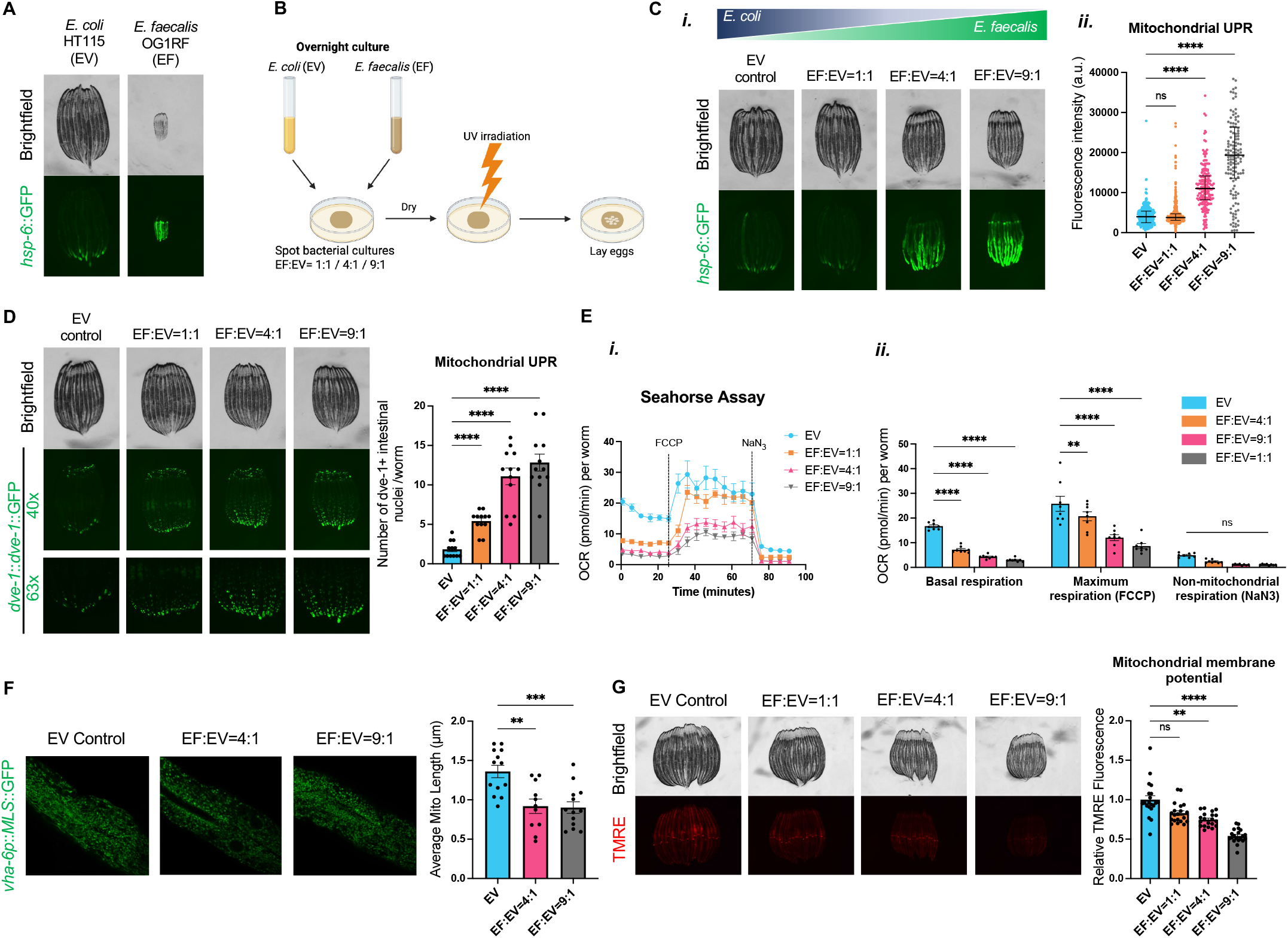
*E. faecalis* infection induces mitochondrial dysfunction and UPR^MT^ activation in C. elegans. (A) Fluorescence images of *hsp-6p*::GFP in D2 adult animals hatched and grown on *E. coli* HT115 or *E. faecalis* OG1RF. (B) Schematic of UPR^MT^ induction assay. (C-G): Mitochondrial assays were conducted on animals grown on mixed cultures of *E. coli* HT115 and *E. faecalis* OG1RF at the indicated ratios. (C) Fluorescence images (left) and quantification (right) of *hsp-6p*::GFP. (D) Fluorescence images (left) and quantification of the total number of intestinal DVE-1 puncta per animal (right) in *dve-1p::dve-1*::GFP animals. (E) Measurement of OCR using a Seahorse XFe96 Analyzer (i) and quantification (ii). (F) Fluorescence images of intestinal mitochondria (left) and quantification of mitochondrial length (right) in *vha-6p*::MLS::GFP animals. *n*=12-15 worms per condition. (G) Fluorescence images (left) and quantification of mitochondrial membrane potential via TMRE staining in N2 animals. Data are represented as median ± interquartile range (C) or mean ± SEM (D/E/F/G). Kruskal-Wallis test with Dunn’s multiple comparisons test (C/G); Brown-Forsythe and Welch ANOVA tests with Dunnett’s T3 multiple comparisons test (D); One-way or two-way ANOVA with Dunnett’s multiple comparisons test (E/F). See also Figures S1 and S2.

We next examined whether this UPR^MT^ activation reflected bona fide mitochondrial damage. Firstly, a Seahorse assay revealed that the oxygen consumption rate (OCR), including both basal and maximal respiration, decreased dose-dependently as the *E. faecalis* ratio increased, indicating impaired ETC activity and systemic bioenergetic failure in *E. faecalis*-exposed animals (Fig. 1E). We then performed confocal imaging of intestine-specific mitochondrial reporter animals (*vha-6p*::MLS::GFP) and assessed mitochondrial morphology. We observed a significant reduction in intestinal mitochondrial length in *E. faecalis*-exposed animals, suggesting an increase in mitochondrial fragmentation (Fig. 1F). Moreover, staining with tetramethylrhodamine ethyl ester (TMRE), a potentiometric dye that accumulates in active mitochondria, demonstrated a dose-dependent loss of mitochondrial membrane potential (Fig. 1G). Together, these data establish that *E. faecalis* causes profound structural and functional damage to host mitochondria, thereby triggering the UPR^MT^ activation. While multiple factors may contribute to the known lethality of *E. faecalis* in *C. elegans,*^19^ our data suggest that mitochondrial dysfunction may be a critical component of this reduced fitness. Indeed, loss of *atfs-1*, a key regulator of the UPR^MT^, exacerbated the shortened lifespan of animals infected with live *E. faecalis*, indicating that the UPR^MT^ serves a partially protective role during *E. faecalis* infection (Fig. S1E).

Having established that *E. faecalis* induces profound and specific insults to host mitochondria, we next sought to identify the underlying cause of the mitochondrial dysfunction. We first asked if this UPR^MT^ activation was a consequence of developmental defects. To determine the temporal requirements for UPR^MT^ activation, we performed transfer experiments by moving animals from *E. coli* to *E. faecalis* at different developmental stages (Fig. S1D-i). We found that larvae remain susceptible to UPR^MT^ induction regardless of the stage of transfer; however, animals transferred to *E. faecalis* as adults failed to activate the response. Additionally, this induction required approximately 48 hours of exposure (Fig. S1D-ii), suggesting that the *E. faecalis*-induced mitochondrial dysfunction results from the cumulative effect of prolonged exposure during development rather than an acute pathogenic insult. This prompted us to investigate whether the UPR^MT^ activation was a general consequence of pathogen-induced developmental defects. To test this, we challenged larvae with *Staphylococcus aureus* and *Staphylococcus saprophyticus*, both of which have been reported to induce severe growth inhibition.^20,21^ Notably, despite inducing comparable levels of developmental arrest, neither *Staphylococcus* strain activated UPR^MT^ (Fig. S2A). This indicates that the UPR^MT^ activation observed in *E. faecalis*-exposed animals is not a secondary effect of growth failure, but a specific response to *E. faecalis*.

Finally, we investigated whether this UPR^MT^ induction requires an active infection. Previous work established that *E. faecalis* pathogenicity requires persistent colonization and proliferation within the *C. elegans* gut lumen, and notably, antibiotic-treated *E. faecalis* was reported to be non-lethal.^19^ However, we found that both heat-inactivated and paraformaldehyde (PFA)-fixed *E. faecalis*, which were confirmed to be metabolically inactive by ATP assay, remained capable of triggering the UPR^MT^ and developmental arrest (Fig. S1F-i, ii). This suggests that the UPR^MT^ activation is not a reaction to active infection or colonization, but is instead triggered by a stable, intrinsic component of *E. faecalis*. Comparing pathogenic isolates (OG1RF and V583) with the probiotic strain DSM16440 further characterized the features of this intrinsic trigger. We found that the probiotic *E. faecalis* strain DSM16440, which lacks the canonical virulence factors associated with clinical infection,^15,22^ induced the UPR^MT^ as robustly as the pathogenic isolates (OG1RF and V583) (Fig. S2B, S2C). Intriguingly, while pathogenic OG1RF caused significantly higher mortality, both the probiotic and pathogenic strains shortened host lifespan (Fig. S2D). This implies that while canonical virulence factors account for the accelerated mortality observed with pathogenic OG1RF, a separate, conserved factor present in both pathogenic and probiotic *E. faecalis* strains drives the UPR^MT^ activation and the baseline reduction in host lifespan.

Taken together, *E. faecalis* exposure induces mitochondrial dysfunction in the host *C. elegans*, which is characterized by impaired mitochondrial respiration, network fragmentation, and membrane potential collapse. Crucially, this response is triggered through a mechanism independent of bacterial viability or canonical virulence factors.

### Nutrient screening identified heme supplementation suppresses *E. faecalis*–induced UPR^MT^

Given that canonical pathogenesis is not required for *E. faecalis*-induced UPR^MT^ activation, we hypothesized the response may be driven by metabolic or nutritional stress. As *C. elegans* depends on the nutrients provided by its bacterial diet, it is plausible that *E. faecalis* is not producing a harmful toxin but instead lacks nutrients vital for host development, thereby inducing mitochondrial stress. To test this hypothesis, we exposed animals to *E. faecalis* mixed with liquid *C. elegans* Maintenance Media (CeMM), an axenic medium composed of various essential vitamins, minerals, and metabolic cofactors that mimic the nutritional components provided by a live *E. coli* diet.^23^ Surprisingly, we found that although live *E. faecalis* supplemented with CeMM still robustly activated the UPR^MT^, CeMM supplementation completely abolished the UPR^MT^ activation induced by PFA-fixed or heat-killed bacteria (Fig. 2A). These results allow us to draw two critical conclusions. First, they confirm that nutritional deficiency is a primary driver of *E. faecalis*-induced UPR^MT^ activation, as providing the missing nutrients via CeMM supplementation was sufficient to turn off the UPR^MT^ induced by dead *E. faecalis*. Second, the failure of CeMM to suppress the UPR^MT^ induced by live *E. faecalis* implies two distinct possibilities: 1) live *E. faecalis* actively scavenges essential nutrients from CeMM and leaves the host nutritionally starved (“Nutrient Competition Model”) or 2) live *E. faecalis* possesses an additional, viability-specific factor that drives mitochondrial stress alongside the nutritional deficiency (“Two-Hit Model”).

**Figure 2.**
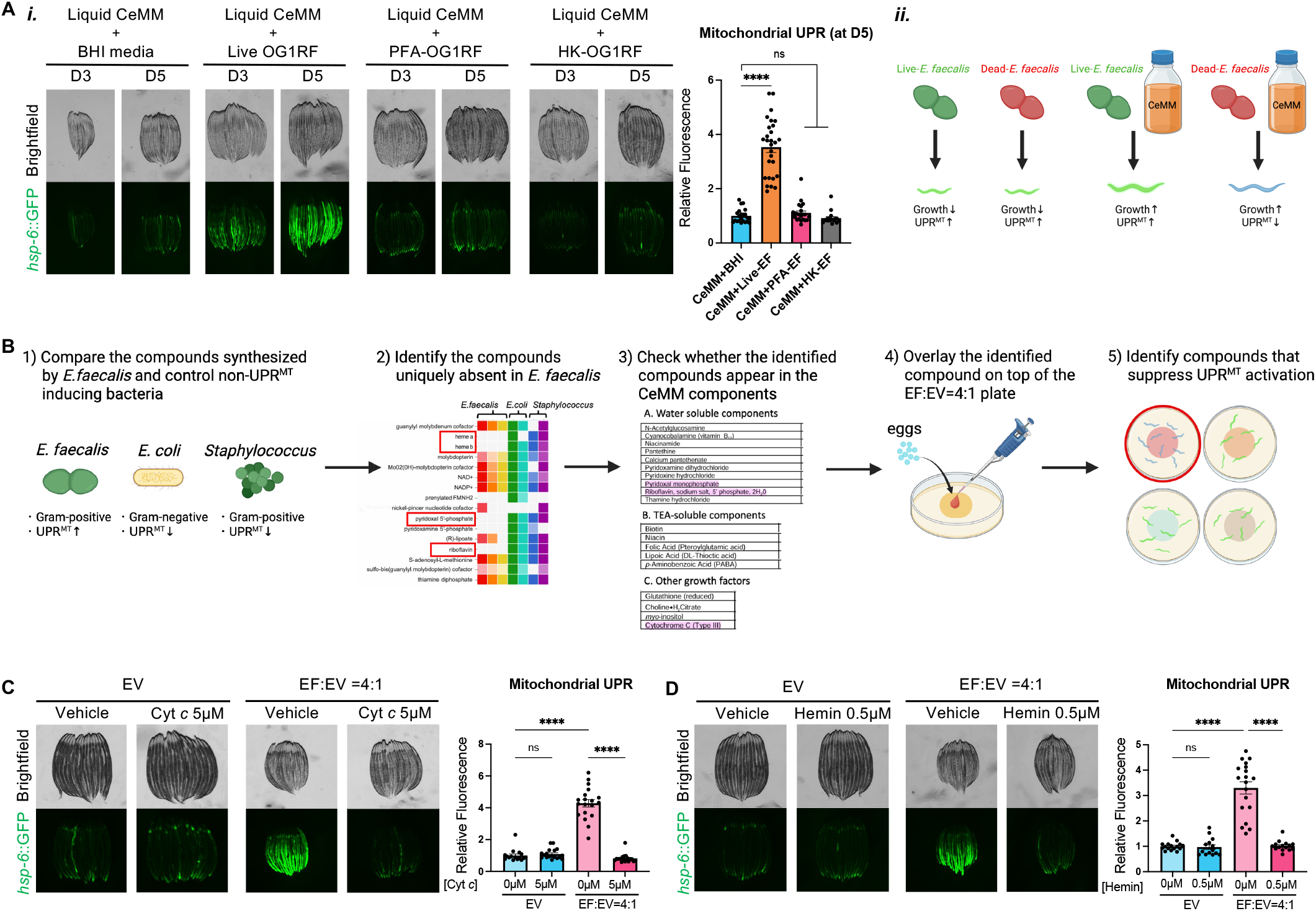
Nutrient screening identified heme supplementation suppresses *E. faecalis*–induced UPR^MT^. (A) Fluorescence images and quantification of *hsp-6p*::GFP in D3 or D5 adult animals hatched and grown in the mixed liquid culture of CeMM and live/dead *E. faecalis* OG1RF (i). Schematic of the results of CeMM supplementation during *E. faecalis* exposure (ii). (B) Schematic of the nutrient screening. (C/D) Fluorescence images (left) and quantification (right) of *hsp-6p*::GFP in D1 adult animals grown on EV bacteria or a 4:1 mixed bacterial plate (*E. faecalis*:EV) supplemented with cytochrome *c* (C) or hemin chloride (D). All data are represented as mean ± SEM. Kruskal-Wallis test with Dunn’s multiple comparisons test (A); Two-way ANOVA with Tukey’s multiple comparisons test (C/D). See also Figure S3.

Here, we first focused on determining the specific dietary factor capable of suppressing the *E. faecalis*-induced UPR^MT^ and conducted a systematic nutrient screen (Fig. 2B). Using the BioCyc database,^24^ we compared the biosynthetic capabilities of *E. faecalis* relative to non-UPR^MT^-inducing control bacteria (*E. coli* as a Gram-negative control and *Staphylococcus* strains as Gram-positive controls) (Fig. S3A). This *in silico* analysis highlighted several bacterial compounds that are uniquely absent in *E. faecalis* but present in the control bacteria (Fig. S3B). We further narrowed down these candidates by cross-referencing them with the CeMM components and supplemented each candidate to *E. faecalis*-exposed animals. Among these candidates, we found that supplementation with cytochrome *c* (Cyt *c*) specifically suppressed the *E. faecalis*-induced UPR^MT^ (Fig. 2C, S3C). Since Cyt *c* is a hemoprotein, we speculated the rescue was driven by its heme cofactor. Consistent with this hypothesis, supplementation with hemin chloride alone similarly abolished the *E. faecalis*-induced UPR^MT^ (Fig. 2D). This rescue is mechanistically consistent with the fact that *C. elegans* is a natural heme auxotroph^25^ and that heme is an essential cofactor for mitochondrial function.^26^ Because host *C. elegans* lacks the biosynthetic machinery to produce heme *de novo* and is entirely dependent on exogenous dietary sources, exposure to *E. faecalis*, which is also a heme-auxotrophic bacterium, likely induced systemic heme starvation, specifically triggering mitochondrial stress. Notably, the supplementation of Cyt *c* or hemin chloride also partially restored the body sizes of *E. faecalis*-exposed animals, implying that heme availability modulates host physiological fitness. Finally, we confirmed that Cyt *c* supplementation failed to suppress the UPR^MT^ induced by *cco-1* RNAi, demonstrating that heme does not globally inhibit mitochondrial stress signaling but specifically rescues mitochondrial stress induced by *E. faecalis* (Fig. S3D).

Collectively, our screening revealed heme as the essential nutrient missing from the *E. faecalis* diet and demonstrated that its supplementation is sufficient to abolish the *E. faecalis*-induced UPR^MT^.

### Heme deficiency is the primary driver of the mitochondrial stress triggered by *E. faecalis*

To confirm that the UPR^MT^ induction is a direct consequence of systemic heme depletion in the host, we next conducted several assays to evaluate the endogenous heme status of *E. faecalis*-exposed animals. We first reanalyzed the published RNA sequencing (RNA-seq) data comparing the L4 larvae transferred and exposed to *E. faecalis* for 16 hours to those exposed to control *E. coli.*^27^ This analysis revealed that the canonical heme-responsive genes (*hrg-1, 2, 3, 4, 7*, and *mrp-5*) are all transcriptionally up-regulated in *E. faecalis*-exposed animals (Fig. 3A). Given that these genes are known to be robustly induced under heme-deficient conditions to maintain host homeostasis,^28,29^ these data suggest that *E. faecalis* exposure results in systemic heme deprivation. Next, to validate these findings in our own experimental system, we performed quantitative polymerase chain reaction (qPCR) analysis on these heme-responsive transcripts. Consistent with the published RNA-seq data, we found that *E. faecalis-*exposed animals significantly up-regulated the expression of these heme starvation markers; however, this transcriptional shift was entirely suppressed by the supplementation with Cyt *c*, confirming that the response is specifically driven by the heme deficiency (Fig. 3B). Next, to confirm this result *in vivo*, we utilized the GFP transcriptional reporter of *hrg-1* as a heme starvation sensor.^30^ Consistent with our qPCR data, we observed a robust induction of *hrg-1p*::GFP expression in *E. faecalis-*exposed animals, which was entirely suppressed by the exogenous heme supplementation. (Fig. 3C). Collectively, these results provide multifaceted evidence that *E. faecalis* exposure causes a severe host heme deficiency.

**Figure 3.**
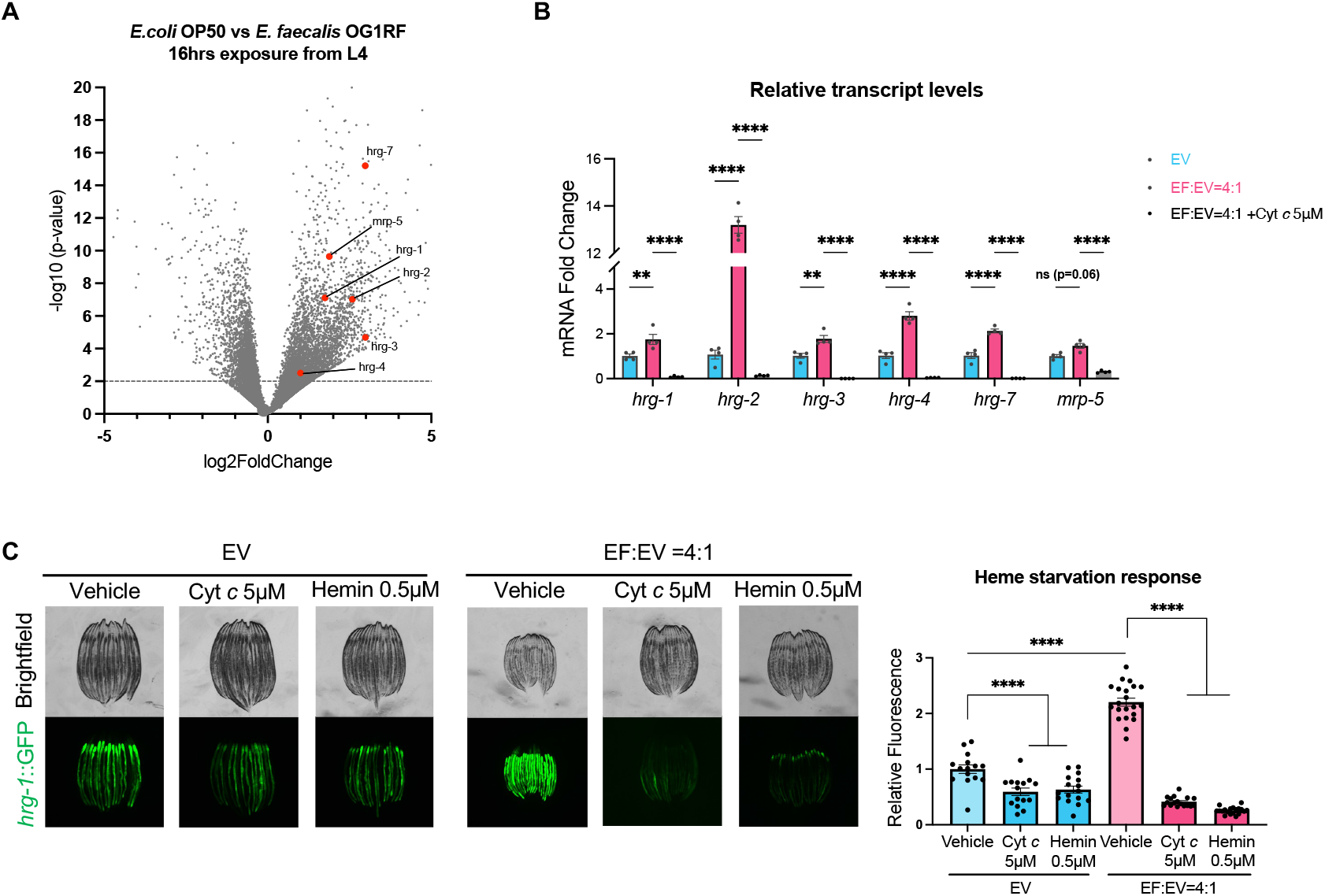
Heme deficiency is the primary driver of the mitochondrial stress triggered by E. faecalis. (A) RNA-seq dataset of wildtype animals exposed to *E. coli* OP50 or infected with *E. faecalis* OG1RF for 16 hours from the L4 stage was reanalyzed for heme response genes. (B) Animals grown on EV bacteria or a 4:1 mixed bacterial plate (*E. faecalis*:EV) with or without cytochrome *c* supplementation were harvested on D1 adult for measuring the expression levels of heme response genes using qPCR. *n*=4 samples per condition. (C) Fluorescence images (left) and quantification (right) of *hrg-1p*::GFP in D1 adult animals grown on EV bacteria or a 4:1 mixed bacterial plate (*E. faecalis*:EV) with or without heme supplementation. Data are represented as mean ± SEM (B/C). Two-way ANOVA with Tukey’s multiple comparisons test (B/C). See also Figure S4.

To determine if heme deficiency alone is sufficient to trigger UPR^MT^ activation independently of other *E. faecalis*-specific factors, we utilized the heme-deficient *E. coli* strain RP523.^31^ This strain lacks the capability to synthesize heme *de novo* due to a mutation in *hemB*, one of the key genes in the heme biosynthetic pathway but remains permeable to exogenous heme; therefore, it is widely used as a known heme auxotroph bacterium.^31^ To test whether dietary heme availability directly regulates UPR^MT^ activation, we titrated hemin chloride into the RP523 diet (0, 0.2, 0.5, and 1 μM). We found that while animals fed RP523 supplemented with low concentrations of hemin chloride (0 and 0.2 μM) exhibited robust induction of both the heme starvation response and the UPR^MT^, supplementation with high concentrations of hemin chloride (1 μM) completely suppressed both responses (Fig. S4A/B). These results demonstrate that the dietary heme deficiency is a conserved trigger for UPR^MT^ activation, rather than a response specific to *E. faecalis* exposure.

Lastly, since *E. faecalis* possesses high-affinity iron sequestration systems,^32^ we asked whether the rescue provided by heme supplementation was a secondary effect of increased general iron availability or was strictly dependent on the heme molecule itself. To test this, we supplemented *E. faecalis*-exposed animals with ferric ammonium citrate (FAC). We found that FAC provided a modest reduction in *E. faecalis*-induced UPR^MT^ activation, although it was insufficient to recapitulate the robust suppression of the UPR^MT^ and the rescue of developmental arrest observed with heme supplementation (Fig. S4C). As *C. elegans* is a natural heme auxotroph that lacks the pathway to synthesize heme from inorganic iron,^25^ this result suggests that *E. faecalis*-exposed animals suffer from a primary deficit of heme rather than iron.

Taken together, our findings demonstrate that dietary heme deficiency is the primary driver of *E. faecalis*-induced UPR^MT^ activation, highlighting a critical role of external heme in maintaining mitochondrial proteostasis in *C. elegans*.

### Heme supplementation rescues mitochondrial dysfunction in *E. faecalis-*exposed C. elegans

As we observed significant mitochondrial impairment upon exposure to *E. faecalis* in *C. elegans* (Fig. 1E/F/G), we next investigated whether heme supplementation could physically and functionally restore mitochondrial homeostasis. Using the nuclear translocation of DVE-1::GFP as an additional indicator of UPR^MT^ activation, we first confirmed that both Cyt *c* and hemin chloride supplementation significantly attenuated *E. faecalis*-induced UPR^MT^. However, this suppression was less pronounced than that observed with the *hsp-6p*::GFP reporter (Fig. 4A) Next, to determine whether the heme supplementation improved organelle physiology, we assessed mitochondrial membrane potential via TMRE staining. We found that supplementation with either Cyt *c* or hemin chloride effectively reversed the mitochondrial depolarization induced by *E. faecalis* (Fig. 4B). Consistent with this, heme supplementation also markedly improved mitochondrial morphology in *E. faecalis*-exposed animals. We found that the severe mitochondrial fragmentation induced by *E. faecalis* was rescued by heme supplementation (Fig. 4C). Finally, to evaluate the functional recovery of the ETC, we measured OCR and found that heme supplementation partially restored OCR in *E. faecalis*-exposed animals (Fig. 4D). Importantly, heme supplementation had no significant effect on mitochondrial membrane potential, morphology, and OCR of *E. coli*-fed control animals, indicating that basal heme levels provided by *E. coli* diet are sufficient for maintaining mitochondrial function under homeostatic conditions. Together, these findings demonstrate that exogenous heme supplementation not only suppresses the UPR^MT^ activation but also fundamentally restores mitochondrial bioenergetics and structural integrity under *E. faecalis* exposure.

**Figure 4.**
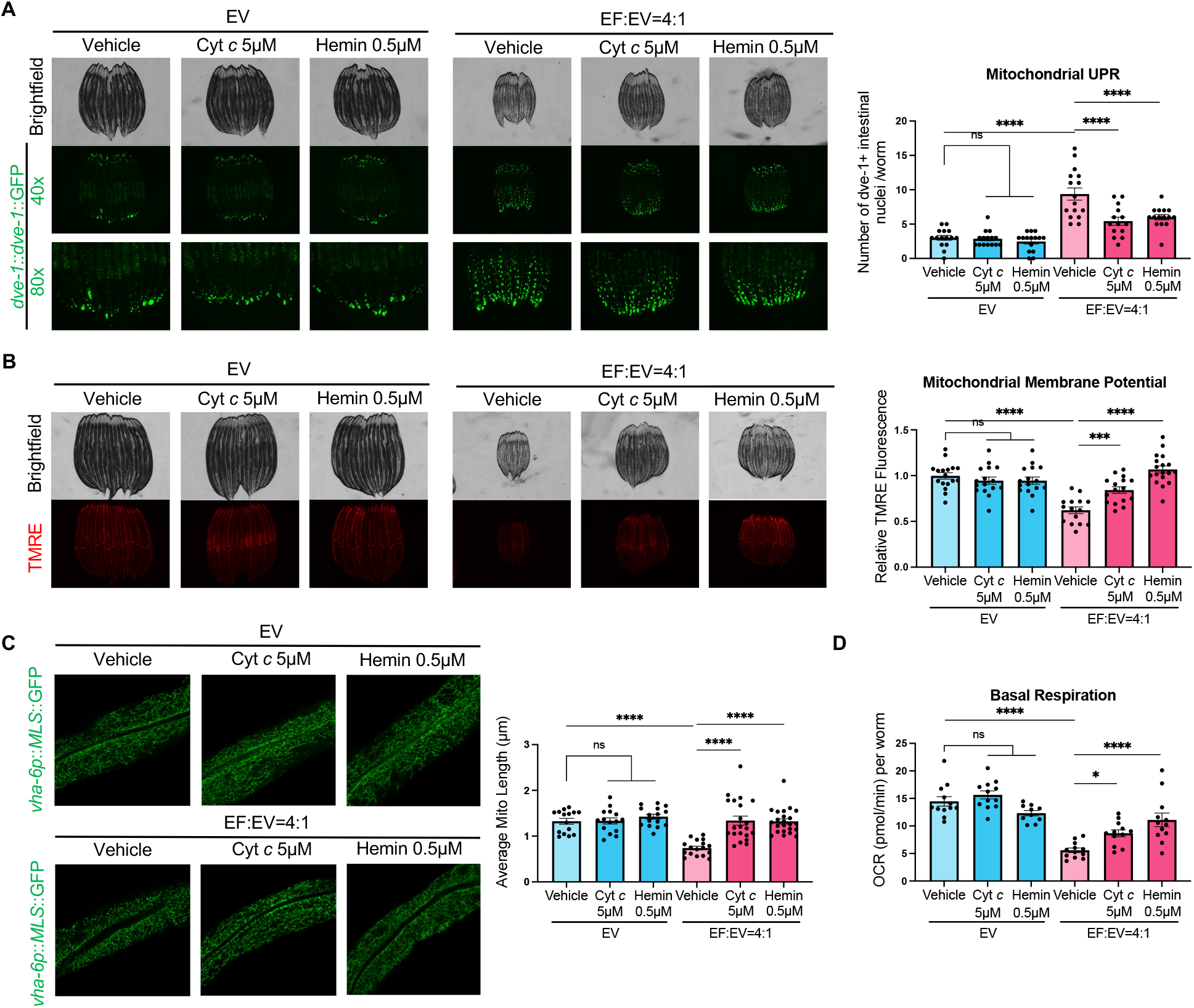
Heme supplementation rescues mitochondrial dysfunction in *E. faecalis-*exposed *C. elegans*. (A–D) Mitochondrial functional assays were conducted in animals grown on EV bacteria or a 4:1 mixed bacterial plate (*E. faecalis*:EV), with or without heme supplementation. (A) Fluorescence images (left) and quantification of the total number of intestinal DVE-1 puncta per animal (right) in *dve-1p::dve-1*::GFP animals. (B) Fluorescence images (left) and quantification of mitochondrial membrane potential (right) via TMRE staining in N2 animals. (C) Fluorescence images of intestinal mitochondria (left) and quantification of mitochondrial length (right) in *vha-6p::MLS*::GFP animals. *n*=15-23 worms per condition. (D) Measurement of basal OCR using a Seahorse XFe96 Analyzer. All data are represented as mean ± SEM. Two-way ANOVA with Tukey’s multiple comparisons test (A/B/C/D).

### Heme supplementation rescues host physiological fitness during *E. faecalis* infection

Having determined that heme supplementation restores mitochondrial integrity, we next asked whether this recovery extends to whole-organism physiology. Given that mitochondria serve as the central hub of energy production, we hypothesized that heme deficiency would create a severe energy deficit, thereby reducing overall host physiological fitness. To test this, we first measured neutral lipid levels using Oil Red O (ORO) staining in animals exposed to *E. faecalis* or *E. coli* controls, with or without heme supplementation. Consistent with a prior report that *E. faecalis* infection induces rapid lipid droplet utilization to fuel the metabolic costs of the immune response,^18^ we observed a significant depletion of lipid content upon *E. faecalis* exposure (Fig. 5A). Notably, this depletion was alleviated by exogenous heme supplementation, suggesting that the severe loss of lipids in *E. faecalis*-exposed animals is driven, at least in part, by heme deficiency and subsequent mitochondrial ETC impairment. Intriguingly, we also found that eggs (F1) laid by *E. faecalis*-exposed parents (P0) exhibited UPR^MT^ activation even prior to hatching (Fig. 5B). In contrast, the P0 generation, which was born from standard *E. coli*-fed animals, only initiated the UPR^MT^ as adults (Fig. 1C). Given that oocytes are heavily provisioned with maternal mitochondria,^33^ it suggests that the F1 generation, born from heme-starved P0 parents, resulted in premature UPR^MT^ activation in eggs due to the lack of maternally inherited heme. Supporting this model, heme supplementation completely abolished the UPR^MT^ activation in offspring laid by *E. faecalis*-exposed animals (Fig. 5B). Additionally, we found that *E. faecalis* exposure significantly reduced the brood size (Fig. 5C). In *C. elegans*, the germline serves as a major sink for metabolic resources, and its activity is tightly coupled to the host’s nutritional and energy status.^34,35^ Thus, this reduced fertility observed in *E. faecalis*-exposed animals likely represents an energy shortage for reproduction or a strategic reallocation of heme for somatic survival. Remarkably, exogenous heme supplementation fully restored reproductive capacity, confirming that heme availability is a primary determinant of germline productivity (Fig. 5C).

**Figure 5.**
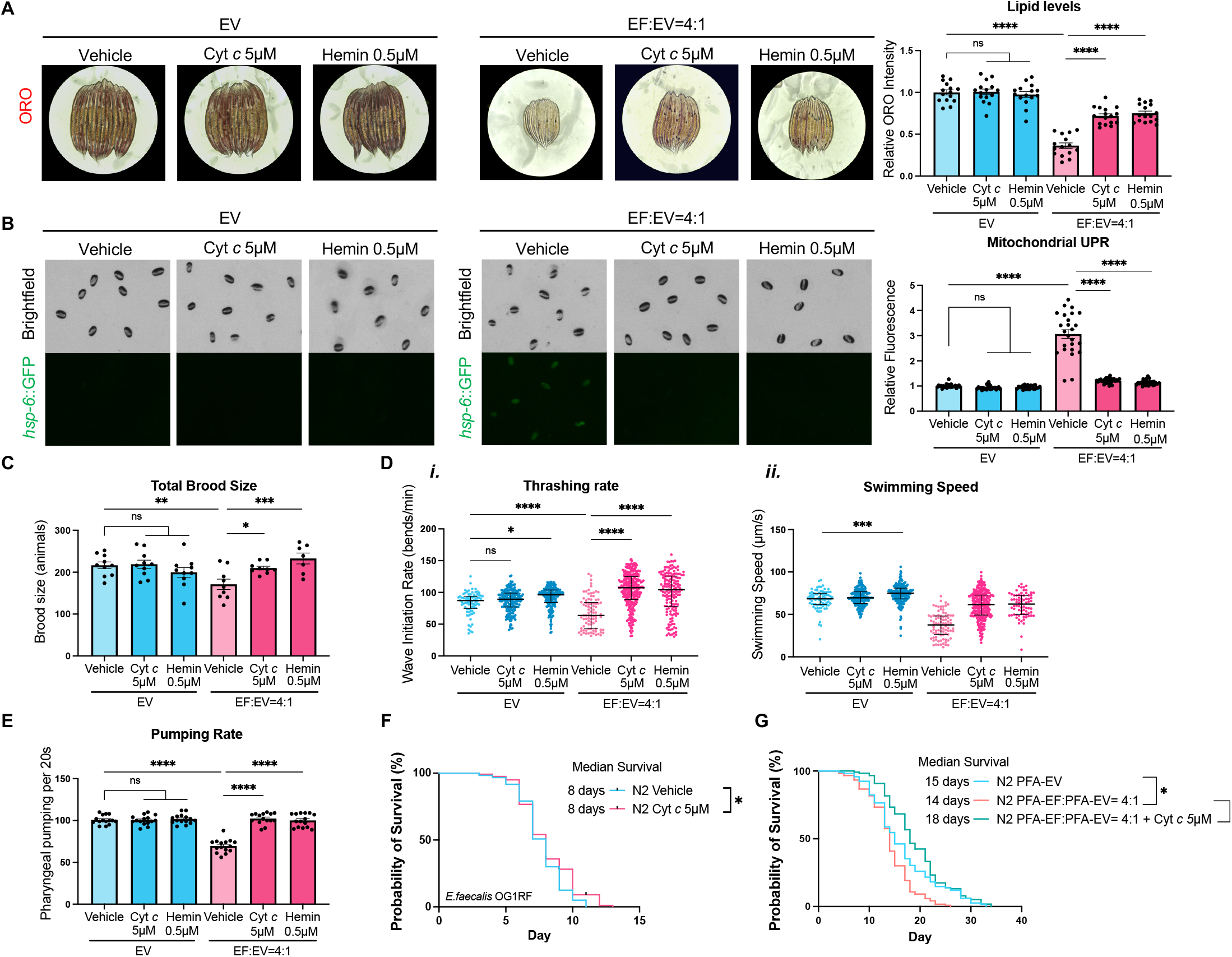
Heme supplementation rescues host physiological fitness during *E. faecalis* infection. (A–E) Physiological fitness of animals grown on EV bacteria or a 4:1 mixed bacterial plate (*E. faecalis*:EV), with or without heme supplementation, was investigated. (A) Oil Red O staining (left) and quantification (right) of lipid levels. (B) Fluorescence images (left) and quantification (right) of *hsp-6p*::GFP in eggs laid by animals grown on the indicated plates. (C) Measurement of total brood size in animals. (D) Measurement of thrashing rate (i) and swimming speed (ii) of animals. (E) Measurement of the pumping rate of animals. (F) Lifespan measurement of animals grown on EV bacteria until the L4 stage, which were then transferred to *E. faecalis* OG1RF mixed with 10% EV bacteria with or without cytochrome *c*. n=120. (G) Lifespan measurement of animals grown on EV bacteria or a 4:1 mixed bacterial plate (*E. faecalis*:EV), with or without cytochrome *c* supplementation. n=120. *N*=2 biological replicates.Data are represented as mean ± SEM (A/B/C/E) or median ± interquartile range (D). Two-way ANOVA with Tukey’s multiple comparisons test (A/B/C/D/E); log-rank test (F/G).

Beyond reproductive fitness, mitochondria are essential for producing the ATP required to maintain physical performance.^36^ To investigate the effects of *E. faecalis*-induced mitochondrial dysfunction on neuromuscular function, we next examined thrashing activity and pharyngeal pumping rates. We found that *E. faecalis* exposure significantly reduced the thrashing activity, as reflected by reduced thrashing rate and swimming speed (Fig. 5D), as well as pharyngeal pumping rate (Fig. 5E). Consistent with our other findings, heme supplementation was sufficient to maintain these rates at control levels, indicating that heme deficiency leads to the reduced energy availability, which was reflected in compromised neuromuscular performance in the host.

Finally, we asked whether these physiological deficits induced by a heme-deficient *E. faecalis* diet contribute to the shortened lifespan of *E. faecalis*-infected animals ^37,38^ (Fig. S2D). First, we supplemented Cyt *c* to animals during live *E. faecalis* infection and observed a slight extension of lifespan. (Fig. 5F). This moderate effect may be explained by the fact that live *E. faecalis* can deploy active virulence factors that damage the host via heme-independent pathways. Moreover, providing heme to live *E. faecalis* may facilitate a metabolic switch from fermentation to respiration and alter their own metabolism,^39,40^ further complicating this host-pathogen interaction. Thus, to limit confounding factors, we next supplemented animals exposed to PFA-killed *E. faecalis* with Cyt *c*. While PFA-killed *E. faecalis* significantly reduced host lifespan compared to PFA-killed *E. coli* control, this defect was remarkably rescued by Cyt *c* supplementation. Notably, the magnitude of this rescue was substantially greater than that observed during live *E. faecalis* infection (Fig. 5G). These results suggest that, while multiple factors contribute to *E. faecalis* pathogenesis, the heme-auxotrophic nature of the bacteria partially explains the shortened lifespan of *E. faecalis*-infected animals. Furthermore, our findings indicate that the loss of dietary heme alone is sufficient to shorten host lifespan.

Collectively, these results identify heme deficiency as a fundamental trigger of mitochondrial dysfunction, ultimately depleting host energy levels and driving systemic physiological decline in *E. faecalis*-exposed animals.

### Live *E. faecalis* infection induces H₂O₂-dependent oxidative stress, leading to UPR^MT^ activation

Our results thus far have established dietary heme deficiency as a primary driver of *E. faecalis*-induced UPR^MT^; however, it remained unclear why nutrient-rich CeMM supplementation completely abolished the UPR^MT^ induced by dead *E. faecalis* but failed to do so during live *E. faecalis* infection (Fig. 2A). This divergence initially pointed to two possibilities: a “Nutrient Competition Model”, where live bacteria scavenge the nutrients from CeMM, or a “Two-Hit Model”, where a second, live *E. faecalis*-specific factor amplifies mitochondrial stress. We first tested the “Nutrient Competition Model” by investigating whether animals supplemented with heme still experienced heme starvation during live *E. faecalis* infection. Remarkably, when we supplemented increasing concentrations of Cyt *c* (5, 10, or 20 μM) with *E. faecalis*-infected animals, it failed to abolish the UPR^MT^ induction even at high concentrations (20 μM) (Fig. 6A), while all three concentrations of Cyt *c* rescued systemic heme deficiency as evidenced by significantly reduced heme-sensitive marker *hrg-1* expression (Fig. S5A). Moreover, a similar result was observed with probiotic *E. faecalis* DSM16440-colonized animals. While Cyt *c* supplementation successfully resolved the systemic heme deficiency (Fig. S5B), it failed to suppress the UPR^MT^ (Fig. S5C). These results effectively rule out the “Nutrient Competition Model” and support the existence of a second layer of stress induced specifically by live *E. faecalis*, independent of classical virulence factors.

**Figure 6.**
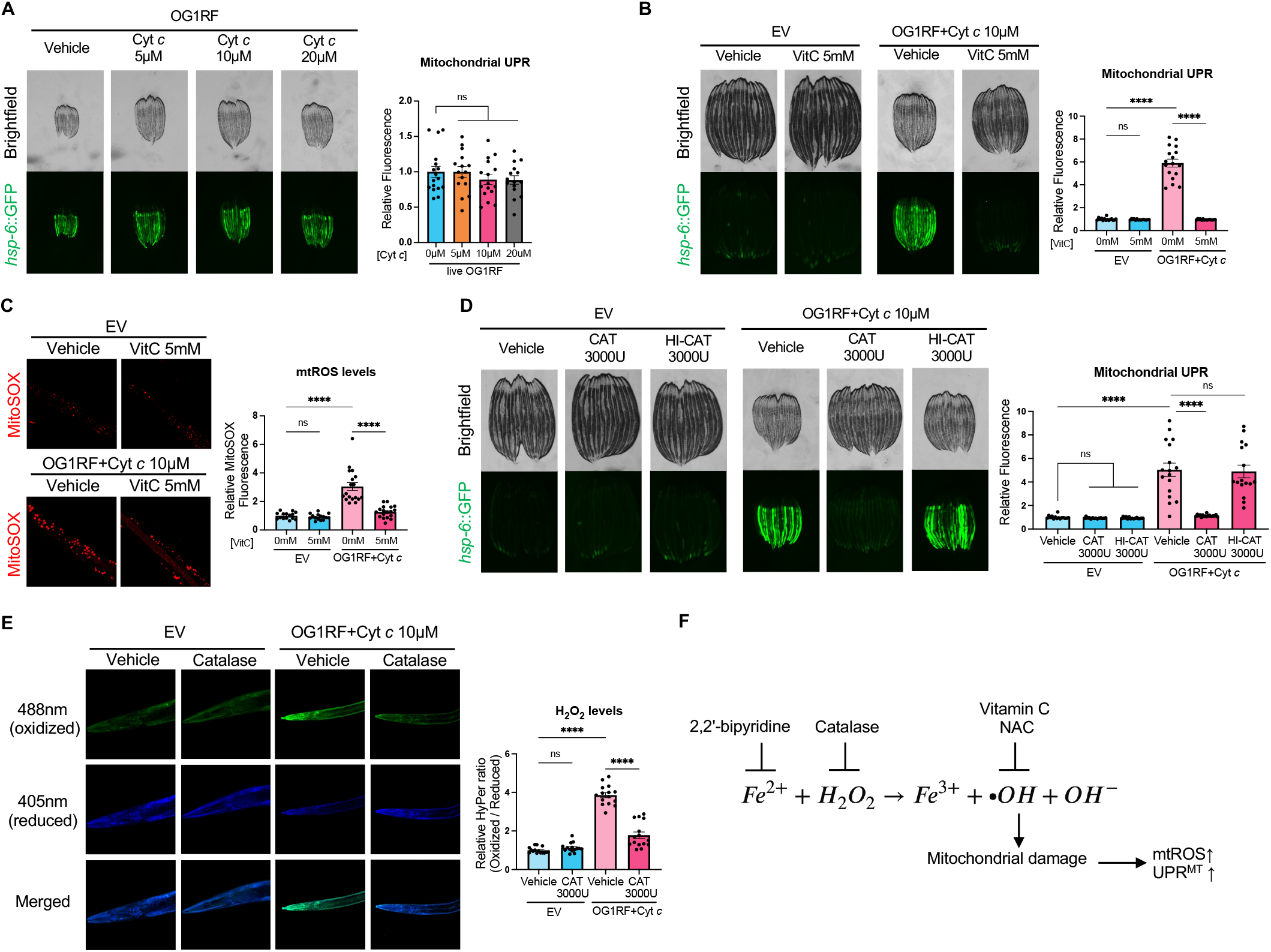
Live *E. faecalis* infection induces H₂O₂-dependent oxidative stress, leading to UPR^MT^ activation. (A) Fluorescence images (left) and quantification (right) of *hsp-6p*::GFP in animals grown on live *E. faecalis* OG1RF supplemented with cytochrome *c* at the indicated concentration. (B/C) Fluorescence images (left) and quantification (right) of *hsp-6p*::GFP (B) or MitoSOX staining (C) in animals grown on live *E. faecalis* OG1RF co-supplemented with cytochrome *c* and vitamin C. (D/E) Fluorescence images (left) and quantification (right) of *hsp-6p*::GFP (D) or Hyper reporter (E) in animals grown on live *E. faecalis* OG1RF co-supplemented with cytochrome *c* and catalase. (F) Schematic of the mechanism of mitochondrial damage induced by the Fenton reaction. All data are represented as mean ± SEM. One-way ANOVA with Dunnett’s multiple comparisons test (A); Two-way ANOVA with Tukey’s multiple comparisons test (B/C/D/E). See also Figure S5.

Live *E. faecalis* infection creates a highly oxidative environment within the host intestinal lumen due to reactive oxygen species (ROS) produced by both bacterial metabolism and host innate immune defenses. On the pathogen side, *E. faecalis* is well documented as a producer of hydrogen peroxide (H_2_O_2_), which acts as a virulence factor against host tissue.^41,42^ Simultaneously, the host *C. elegans* utilizes the dual oxidase BLI-3, which localizes to the intestinal cell membrane, to secrete H_2_O_2_ into the gut lumen to neutralize pathogens and enhance immunity.^42,44,45,45^ Given these parallel sources of ROS, we hypothesized that H_2_O_2_, generated by either the host or bacteria, may serve as an additional stressor on top of heme deficiency and drive mitochondrial damage during active infection. To test this, we co-supplemented the antioxidant Vitamin C (VitC) alongside Cyt *c* to live *E. faecalis*-infected animals. Supporting our hypothesis, we found that the VitC co-supplementation completely abolished the UPR^MT^ induction observed when Cyt *c* alone was supplemented to *E. faecalis*-infected animals (Fig. 6B). Consistent with this, we confirmed that *E. faecalis* infection significantly induced mitochondrial superoxide levels, as measured by MitoSOX fluorescence, which was suppressed by VitC co-supplementation (Fig. 6C). Additionally, this protective effect of an antioxidant was similarly observed with another broad ROS scavenger, N-acetylcysteine (NAC) (Fig. S5D). However, co-supplementation with TEMPOL, a superoxide dismutase mimetic that rapidly converts superoxide into H_2_O_2_, failed to suppress the UPR^MT^ (Fig. S5E), suggesting that H_2_O_2_ rather than upstream superoxide was the specific driver of the mitochondrial damage and subsequent UPR^MT^ activation.

To directly test the involvement of H_2_O_2_, we next co-supplemented *E. faecalis*-infected animals with catalase (CAT), an enzyme that neutralizes H_2_O_2_ by breaking it down into water and oxygen. Consistent with our hypothesis, CAT supplementation effectively suppressed the UPR^MT^ activation observed in *E. faecalis*-infected animals supplemented with Cyt *c* alone (Fig. 6D). Importantly, this rescue was CAT dose-dependent (Fig. S5F), and required enzymatic activity, as heat-inactivated catalase (HI-CAT) failed to suppress the stress response (Fig.6D). Furthermore, measurement of intracellular H_2_O_2_ levels using Hyper fluorescent reporter (*rpl-17p*::HyPer) revealed a significant increase in cytosolic H_2_O_2_ in *E. faecalis*-infected animals, which was significantly attenuated by CAT supplementation (Fig. 6E). Together, these data demonstrate that extracellularly generated H_2_O_2_ can diffuse into host intestinal cells to act as a critical mediator of the live *E. faecalis*-specific mitochondrial stress.

We next asked how H_2_O_2_ specifically induced mitochondrial stress during *E. faecalis* infection. While H_2_O_2_ is relatively stable, non-radical ROS, it exerts severe cellular toxicity when it reacts with intracellular labile iron to generate highly reactive free hydroxyl radicals (·OH) via the Fenton reaction.^47,48,49^ We reasoned that if the Fenton reaction drives this secondary mitochondrial stress, sequestering the available ferrous iron (Fe^2+^) should prevent the reaction and protect the mitochondria. Strikingly, supplementation with the cell-permeable iron chelator 2,2’-bipyridine successfully suppressed the UPR^MT^ activation observed during *E. faecalis* infection (Fig. S5G). As mitochondria are highly enriched in iron, these results suggest they become a major target for this Fenton-mediated damage. Together, these findings establish the Fenton reaction as the critical secondary stressor that exacerbates mitochondrial damage during live *E. faecalis* infection (Fig. 6F).

### The host dual oxidase BLI-3 is the primary source of H₂O₂ fueling mitochondrial dysfunction during *E. faecalis* infection

Lastly, we sought to determine whether the H_2_O_2_ driving this secondary stress originates from the bacteria or the host. We first asked whether bacterial metabolism was the primary source of H_2_O_2_. To test this, we utilized *E. faecalis* transposon mutants of H_2_O_2_-production pathways, including glycerol-3-phosphate oxidase (*glpO*), NADH oxidase (*nox*), and the extracellular electron transfer (EET) pathways.^50,51^ However, animals infected with these mutants maintained UPR^MT^ activation indistinguishable from that of the wild-type control (Fig. S6A) despite reduced H_2_O_2_ production observed in EET pathway mutants (Fig. S6B). Having ruled out pathogen-derived H_2_O_2_ as the driver of the Fenton reaction, we next tested whether host BLI-3-derived H_2_O_2_ is responsible for triggering the secondary mitochondrial stress. To test this, we evaluated UPR^MT^ activation in *bli-3(im10)* mutants, which lack a functional oxidase domain and are known to generate reduced amounts of H_2_O_2_ upon *E. faecalis* infection.^46^ Remarkably, the *bli-3(im10)* mutant completely abolished the UPR^MT^ activation during *E. faecalis* infection (Fig. 7A). Additionally, we utilized *tsp-15(sv15)* mutants, which lack the functional tetraspanin protein required for the functional assembly and H_2_O_2_-generating activity of the BLI-3 complex at the apical membrane.^52,53^ Consistent with the *bli-3 (im10)* mutants, the loss of TSP-15 also led to the marked suppression of UPR^MT^ activation (Fig. 7A). Importantly, both *bli-3(im10)* and *tsp-15(sv15)* mutants exhibited robust UPR^MT^ activation upon exposure to the canonical UPR^MT^ inducer *cco-1* RNAi, confirming that these animals retain a fully intact UPR^MT^ machinery (Fig. 7B). Furthermore, we observed a significantly attenuated mtROS level in *bli-3(im10)* animals, indicating that BLI-3-derived H_2_O_2_ is the underlying cause of mitochondrial oxidative damage (Fig. 7C). Together, these results demonstrate that the host dual oxidase BLI-3, rather than bacterial metabolism, is the primary source of the H_2_O_2_ that fuels the Fenton reaction and unintentionally drives the secondary mitochondrial stress during *E. faecalis* infection.

**Figure 7.**
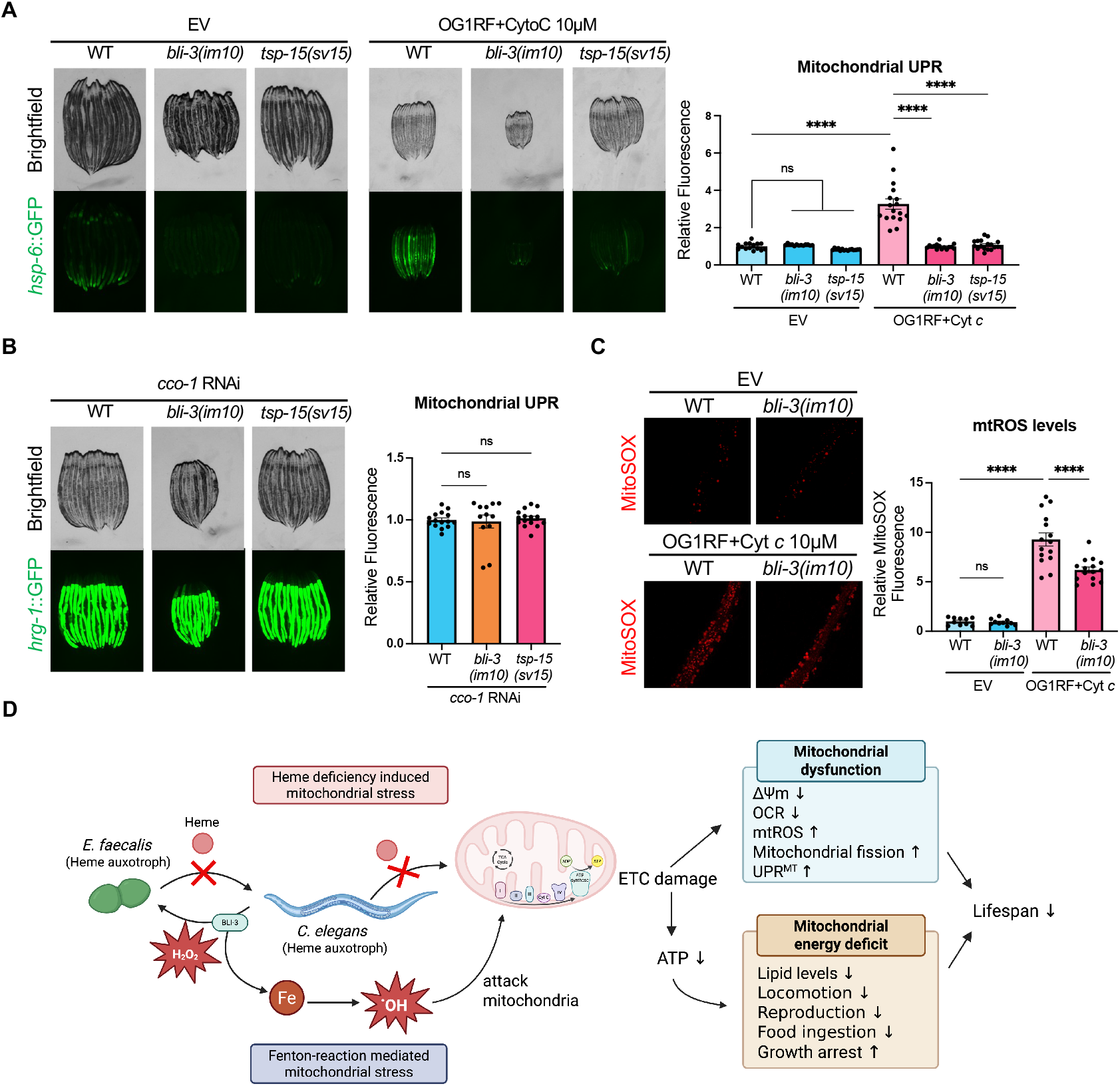
The host dual oxidase BLI-3 is the primary source of H₂O₂ fueling mitochondrial dysfunction during *E. faecalis* infection. (A) Fluorescence images (left) and quantification (right) of *hsp-6p*::GFP in wildtype, *bli-3(im10), tsp-15(sv15)* animals grown on live *E. faecalis* OG1RF supplemented with cytochrome *c*. (B) Fluorescence images (left) and quantification (right) of *hsp-6p*::GFP in wildtype, *bli-3(im10), tsp-15(sv15)* animals grown on *cco-1* RNAi bacteria. (C) Fluorescence images (left) and quantification (right) of MitoSOX staining in wildtype or *bli-3(im10)* animals grown on live *E. faecalis* OG1RF supplemented with cytochrome *c*. *N*=2 biological replicates. (D) Schematic of the UPR^MT^ induction by *E. faecalis* infection. All data are represented as mean ± SEM. Two-way ANOVA with Tukey’s multiple comparisons test (A/C); Kruskal-Wallis test with Dunn’s multiple comparisons test (B). See also Figure S6.

## Discussion

In this study, we have identified a synergistic mechanism by which *E. faecalis*-induced mitochondrial proteotoxic stress leads to UPR^MT^ activation in the host *C. elegans*. In this “Two-Hit Model”, we demonstrate that this mitochondrial dysfunction observed during *E. faecalis* infection is driven by the combination of bacteria-induced host heme starvation and a host-derived immune response (Fig. 7D). Our discovery provides a mechanistic explanation for how a luminal pathogen can trigger mitochondria-specific damage without the direct secretion of mitochondrial toxins, while presenting a new perspective that UPR^MT^ activation functions as a critical homeostatic sensor for metabolic imbalances during infection.

A primary finding of our work is the role of heme as a vital nutrient required for both mitochondrial and organismal health. While heme is universally recognized as a fundamental cofactor, its precise role in maintaining host homeostasis is difficult to isolate in mammalian models due to confounding factors arising from endogenous heme synthesis pathways. In contrast, *C. elegans* is a natural heme auxotroph, making it a uniquely tractable *in vivo* model for investigating the organellar and physiological consequences of heme deficiency.^25^ In our study, this *C. elegans-E. faecalis* interaction model unexpectedly created a heme-deficient state in the host due to the heme-auxotrophic nature of both bacteria and the host, leading to the discovery of heme starvation-derived UPR^MT^ activation. Notably, a previous large-scale screen of bacterial species isolated from the natural habitat of *C. elegans* found that approximately 18% of over 500 bacterial strains activated UPR^MT^.^54^ Our findings imply the intriguing possibility that some of these previously identified commensal or pathogenic bacteria may also activate the UPR^MT^ in a virulence factor-independent manner by altering environmental nutrient availability.

A second highlight of our work is the identification of a “double-edged sword” in the host innate immune response. The BLI-3/TSP-15 dual oxidase complex is well characterized as a frontline defense mechanism that neutralizes luminal pathogens by secreting H_2_O_2.43_^,44,45,46,52,53^ Consistent with their roles, the BLI-3/TSP-15 complex is critical for surviving pathogen encounters.^45,46^ However, this host-derived H_2_O_2_ cannot specifically target *E. faecalis* and instead diffuses back into the host intestinal tissue, where it reacts with labile iron to trigger the Fenton reaction in iron-rich mitochondria. The resulting hydroxyl radicals immediately damage the host’s mitochondria, demonstrating that the host’s innate immune system can become a driver of its own pathology. Thus, the UPR^MT^ activation observed during *E. faecalis* infection is largely a signature of immunopathology, an energetic cost paid by the host due to a hyperactive immune response. Intriguingly, while we observed a reduced UPR^MT^ activation in *bli-3(im10)* and *tsp-15(sv-5)* animals during *E. faecalis* infection (Fig. 7A), previous studies have shown that *bli-3* KO results in a shortened lifespan during *E. faecalis* infection,^45,46^ suggesting that the bactericidal effect of BLI-3-generated H_2_O_2_ is still stronger than the unwanted secondary action on the host. Moreover, in addition to mitochondria, lysosomes are also known as major hubs for iron storage,^55,56^ suggesting the possibility that *E. faecalis* infection and the subsequent Fenton-mediated oxidative burst may also affect lysosomal function.

Another interesting observation is that the UPR^MT^ was also activated in animals exposed to the probiotic *E. faecalis* DSM16440 (Fig. S5B). This is particularly notable as DSM16440 lacks canonical virulence factors of *E. faecalis* and is used as a probiotic in humans. Although further mechanistic research is required to test the conservation of this model in mammalian systems, our results indicate that even probiotic bacterial colonization may impose a previously unrecognized oxidative stress on the host by triggering the host’s immune response to generate H_2_O_2._ Given that the human microbiome comprises over 3,000 bacterial species,^57,58^ even commensal microbes might occasionally disrupt host metabolic homeostasis by activating host immune surveillance mechanisms.

Collectively, these findings highlight a new conceptual framework for host-pathogen interactions. Pathogen-induced mitochondrial dysfunction has traditionally been interpreted as a direct consequence of bacterial toxicity and specialized virulence factors.^59,60^ However, our findings provide a novel scenario where mitochondrial stress arises from host-pathogen metabolic imbalance and the immunopathological cost of host immune response. Moreover, the evolutionary conservation of these pathways suggests that this mechanism might have broad implications for mammalian pathogenesis. First, while mammals possess endogenous heme synthesis pathways, severe infections may induce a functional heme-deficient state via a host immune response known as “nutritional immunity.” During this response, the host sequesters circulating heme and iron, thereby restricting pathogen access to these trace metals. While it is a defense strategy against bacterial pathogens, it can inadvertently disrupt host metabolism and induce heme deficiency, leading to the clinical condition known as anemia of inflammation (AI) or anemia of chronic disease (ACD).^61,62^ Thus, our findings raise the possibility that the heme deficiency-induced UPR^MT^ observed in *C. elegans* may also occur in mammals as a secondary consequence of nutritional immunity. Second, the mammalian orthologs of BLI-3 are DUOX1 and DUOX2,^44,45,63^ which are primary source of ROS in the human gut and is frequently up-regulated during dysbiosis and inflammatory bowel disease (IBD). Given that *E. faecalis* is a major cause of these diseases, our findings may explain the localized mitochondrial collapse seen during *E. faecalis* induced sepsis or chronic inflammation.^64^ Ultimately, understanding these self-inflicted side effects of the host’s innate immunity may provide a new therapeutic framework for mitigating tissue damage.

### Limitations of the Study

This study used *C. elegans* as a model and identified heme-deficiency-induced UPR^MT^, but the severe heme starvation induced by *E. faecalis* feeding is fundamentally linked to the nematode’s natural heme auxotrophy. Whereas *E. faecalis* infection may also induce heme deficiency in mammals via the activation of nutritional immunity, further studies are required to test this possibility in mammalian models. Similarly, it remains unclear whether *E. faecalis* infection triggers DUOX1/DUOX2-mediated oxidative stress and mitochondrial damage in mammals. Additionally, this study could not directly measure the Fenton reaction within mitochondria; instead, we relied on secondary physiological readouts, such as mtROS levels and UPR^MT^ activation, as indicators of this localized oxidative damage. Finally, given that the mammalian gut is colonized by thousands of bacterial species with diverse metabolic requirements, whether the specific immunopathology observed in this study manifests within a more complex polymicrobial environment remains an important question for future research.

## ACKNOWLEDGMENTS

We thank all members of the Dillin lab for discussion and technical assistance. This work was supported by the Howard Hughes Medical Institute (L.J., A.D.); Ito Foundation Fellowship (A.M.); QUAD Fellowship (A.M.); JASSO Graduate Fellowship for Degree Seeking Students (A.M.); Larry L. Hillblom Foundation Postdoctoral Fellowship (H.Z.); Glenn Foundation for Medical Research Postdoctoral Fellowship (H.Z.).

## AUTHOR CONTRIBUTIONS

Conceptualization: A.M. and A.D. Methodology: A.M. and H.Z. Investigation: A.M., H.Z., A.M.V., and L.K.J. Formal analysis: A.M. Writing - Original Draft: A.M. Writing - Review & Editing: A.M., H.Z., and A.D. Visualization: A.M. Funding Acquisition: A.D. and A.M. Supervision: A.D.

## STAR METHODS

### RESOURCE AVAILABILITY

#### Lead Contact

Further information and requests for resources and reagents should be directed to and will be fulfilled by the lead contact, Dr. Andrew Dillin.

#### Materials Availability

All *C. elegans* strains and bacterial strains used in this study are available by direct request to the lead contact, Dr. Andrew Dillin.

#### Data and Code Availability

All data reported in this paper will be shared by the lead contact upon request.

## EXPERIMENTAL MODEL AND STUDY PARTICIPANT DETAILS

### *C. elegans* maintenance and bleach synchronization

All *C. elegans* strains used for research are variations of N2 sourced from the Caenorhabditis Genetics Center (CGC). For stock maintenance, *C. elegans* were cultured on standard nematode growth media (NGM) agar plates seeded with *Escherichia coli* strain OP50 at 15 °C. All the experiments were performed using bleach synchronized hermaphroditic worms at 20 °C unless otherwise stated. Bleach synchronization was achieved by collecting gravid adults grown on OP50 with M9 solution (22 mM KH_2_PO_4_ monobasic, 42.3 mM Na_2_HPO_4_, 85.6 mM NaCl, and 1 mM MgSO_4_) and incubating for 5 minutes in a standard bleaching solution (1.8% sodium hypochlorite, 0.375 M KOH) to dissolve adults and keep only eggs. Intact eggs were then washed three times with M9 solution and plated. For RNAi experiments, *C. elegans* were fed *Escherichia coli* HT115 with pL4440 plasmids expressing dsRNA targeted to the gene of interest or an empty vector (EV) control and grown on NGM plate containing 1 mM IPTG and 100 mg/mL carbenicillin. All RNAi vectors were obtained from the Vidal library or the Ahringer library and sequence verified. As WT animals, fluorescent reporter alone in an N2 genetic background were used for imaging experiments, and N2 strain were used for all other experiments unless otherwise stated. The IQ6011 strain (*hrg-1p*::GFP) was a kind gift from Dr. Iqbal Hamza (The University of Maryland). The *bli-3(im10)* was a kind gift from Dr. Andrew Chisholm (University of California, San Diego) and Dr. Hiroki Moribe (Kurume University).

## METHOD DETAILS

### Bacterial culturing

Bacterial liquid cultures were prepared prior to the *C. elegans* bacterial exposure assay. *Enterococcus faecalis* strains (OG1RF, V583, DSM16440) were inoculated in brain heart infusion (BHI) broth and grown overnight at 37 °C. The overnight culture was spread on either BHI agar plate or NGM agar plate depending on the experiment.

*Staphylococcus aureus* SA113 and *Staphylococcus saprophyticus* strain NCTC 7292 were cultured in LB broth overnight at 37 °C. 200 uL of overnight culture was spread on 6 cm Nematode Growth Medium (NGM) plates and grown overnight at 37 °C.

### UPR^MT^ induction assay using *E. faecalis*

For the UPR^MT^ induction assay with *E. faecalis*, the OD600 of the bacterial cultures was measured, and the colony-forming units (CFU) were adjusted to achieve an *E. faecalis to E. coli* ratio of 4:1. Specifically, each NGM plate was seeded with 40.5×10^7^ CFU of *E. faecalis* and 12.5×10^7^ CFU of *E. coli* HT115, yielding a total of 53×10^7^ CFU. After drying at room temperature, the plates seeded with the mixed bacterial culture were exposed to UV for 5 min using a UV crosslinker (CL-1000, UVP) to fix this bacterial ratio. This *E. faecalis and E. coli* mixed-feeding condition was utilized to alleviate the growth-arrested phenotype induced by a 100% *E. faecalis* diet, while maintaining robust UPR^MT^ activation.

To activate the UPR^MT^ with metabolically active live *E. faecalis*, animals were hatched on NGM plates seeded with *E. coli* HT115 and transferred to BHI plates seeded with *E. faecalis* at different developmental stages.

### Pharmacological Treatment in *C. elegans*

Stock solutions of cytochrome *c* (C2506, Sigma-Aldrich), Vitamin C (A4544, Sigma-Aldrich),, FAC (F5879, Sigma-Aldrich), NAC (A7250, Sigma-Aldrich), TEMPOL (176141, Sigma-Aldrich), and catalase (C1345, Sigma-Aldrich), were prepared in ultrapure water (Mili-Q, Millipore) at concentrations of 2 mM, 500 mM, 500 mM, 200 mM, 500 mM, and 30,000 U/mL, respectively. Each stock solution was filtered through a 0.2 µm filter and stored at −20 °C. The drugs were overlaid on top of the bacterial lawn to achieve the final concentrations indicated in the figures, and the plates were allowed to dry completely at room temperature before seeding eggs. For heat-inactivated catalase (HI-CAT) supplementation, the filtered catalase stock solution was heated at 80 °C for 10 minutes.

Hemin chloride (3741, Sigma-Aldrich) was dissolved in DMSO at a concentration of 1 mM, and 2,2′-Bipyridine (D216305, Sigma-Aldrich) was dissolved in ethanol at a concentration of 5 mM. These stock solutions were filtered through a 0.2 µm filter. The 1 mM hemin chloride solution was added to the melted NGM agar medium after autoclaving at a final concentration of 0.5 µM. The 5mM 2,2′-Bipyridine stock was diluted 1:10 with ultrapure water (Mili-Q, Millipore) to make a working stock of 500 µM, which was subsequently overlaid on the bacterial lawn.

### Heme limitation assay using *Escherichia coli* RP523

To control the amount of heme supplemented to worms, the heme-deficient *Escherichia coli* strain RP523.^31^ was used, adapting a previously published method.^65^ Briefly, *E. coli* RP523 was grown overnight in LB broth supplemented with 1 µM of hemin chloride. On the following day, the overnight culture was diluted at a ratio of 1:50 into fresh LB broth supplemented with 0, 0.2, 0.5, or 1 μM of hemin chloride and grown overnight. RP523 supplemented with hemin chloride was washed twice with M9 to remove the excess free hemin from media, and 200 uL of the bacterial culture was spread on 6 cm NGM plates.

### Heat-killed and PFA-killed bacterial preparation

Dead bacteria were prepared using either heat-killing or PFA-killing methods, following previously published protocols.^20,66^ For heat-killed bacterial preparation, an overnight bacterial culture was heated at 80 °C for 2 h using a thermomixer (Eppendorf). For PFA-killed bacterial preparation, PFA was added to an overnight bacterial culture to a final concentration of 0.5%, and the culture was incubated at 37 °C for 1 h to metabolically inactivate the bacteria, followed by five washes with fresh LB. The complete killing of bacteria was confirmed by streaking the treated bacteria onto an LB plate and verifying the absence of colony growth.

### Nutrient screening using CeMM

The chemically defined *C. elegans* maintenance medium (CeMM) was prepared according to the formulation previously described.^23,67^ For supplementing CeMM to *E. faecalis*-exposed animals, bleach-synchronized eggs were grown in a solution consisting of 2.5 mL of 2× CeMM and 2.5 mL of bacterial culture (BHI media control, PFA-treated *E. faecalis*, HK-*E. faecalis*, or live *E. faecalis*) in a 15 mL Falcon tube. The tubes were incubated at 20 °C while rotating, and animals were imaged on Day 3 and Day 5 adults.

For the nutrient screening, candidate CeMM components were identified through comparative biology approach as described in the main text. The selected individual CeMM components were added to a 4:1 mixed bacterial plate (*E. faecalis:E. coli*) at the concentrations used for standard CeMM.

### Fluorescence stereomicroscope imaging and quantification in *C. elegans*

To measure the expression of fluorescent reporters in *C. elegans*, animals were randomly picked under a nonfluorescent stereomicroscope and anesthetized using a 100 mM sodium azide solution on NGM plates. Immobilized worms were aligned head-to-tail with a worm pick and imaged with a Leica M250FA automated fluorescence stereomicroscope equipped with a Hamamatsu ORCA-ER camera. For imaging experiments using the *hsp-16.2p*::GFP reporter strain, Day 1 animals were heat-shocked for 2 h at 34 °C and imaged on Day 2.

Quantification of the intestinal GFP signal was performed with a worm sorter (350-5000-000, Union Biometrica) or ImageJ/FIJI. For quantification using a worm sorter, the remaining population was washed off and run through the worm sorter on the same day after imaging, and all fluorescence values (*hsp-6p*::GFP) obtained from the worm sorter were normalized to extinction values (EXT, a measure of optical density used as an indicator of worm size). For quantification using ImageJ/FIJI, intestinal regions of each worm were traced, and the mean intensity was measured. For quantification of the *dve-1p*::DVE-1::GFP reporter strain, the total number of intestinal DVE-1 puncta per worm was manually counted from 63× images.

### Quantitative RT-PCR

Synchronized animals grown on respective bacterial diets were collected on Day 2 of adulthood and gravity washed in M9 buffer to isolate adults and remove larvae. Four biological replicates were prepared per condition, and each replicate contained about 1500 animals. Washed animals were then resuspended in 1 mL of TRIzol (15596026, Invitrogen) and immediately frozen in liquid nitrogen. RNA was harvested by repeating freeze-thaw cycles for three times and following a TRIzol-based extraction method as previously described.^68^ RNA was purified using a RNeasy Mini Kit (74106, Qiagen), and cDNA was synthesized using the QuantiTect Reverse Transcription Kit (205314, Qiagen) with 1 μg of RNA per sample. qPCR was performed using a standard curve protocol using SYBR Select Master Mix (4472920, Applied Biosystems).

### Quantification of mitochondrial features using FIJI

The mitochondrial length of worms was measured following a previously published protocol using intestinal mitochondrial GFP reporter worms (*vha-6p*::MLS::GFP(65C)::*unc-54* 3’UTR). ^69^ Briefly, animals were lightly immobilized using a 10 mM sodium azide solution on a glass slide and immediately imaged using a Zeiss LSM900 Airyscan super-resolution confocal microscope to acquire Z stacks.

For quantification using ImageJ/FIJI, Z-stack fluorescence images were projected into a single layer using the “Average Intensity” function. To isolate the mitochondrial signal, images were then segmented using the “Auto Local Threshold” function (parameters: method=Bernsen, radius=15, parameter_1=0, parameter_2=0, white). Individual mitochondria were then structurally delineated using the “Skeletonize (2D/3D)” function, and the “Analyze Skeleton (2D/3D)” function was applied to measure the length and associated morphological parameters of each mitochondrion.

### TMRE staining

*C. elegans* were stained with TMRE (T669, Thermo-Fisher) as previously described.^70^ In short, Day 1 adult animals were transferred to a corresponding plate overlayed with 100 nM TMRE and incubated overnight at 20 °C shielded from light. On the following day, animals were destained of excess TMRE dye by transferring onto to OP50-seeded NGM plate for 1 h prior to imaging. Images were taken using Leica M250FA automated fluorescence stereomicroscope equipped with a Hamamatsu ORCA-ER camera. Quantification was performed using ImageJ/FIJI software by tracing intestinal regions and measuring the mean intensity.

### Mitochondrial oxygen consumption rate assay

Measurement of the oxygen consumption rate (OCR) was performed using a Seahorse XFe96 Analyzer (Agilent) at 20 °C following a previously published protocol.^69^ Briefly, the sensor cartridge, hydrated overnight in calibrant buffer at room temperature, was loaded with drugs to achieve final well concentrations of 10 µM FCCP and 40 mM NaN_3_ and then calibrated. Animals grown on the respective bacterial diets, with or without heme supplementation, were washed with M9 buffer, and 10–20 animals per well were loaded into a Seahorse XFe96 Cell Culture Microplate (34222, Agilent) for OCR measurement. After completion of the assay, the exact number of animals in each well was counted and OCR values were normalized per worm. To account for differences in worm size among groups, the per-worm OCR was further normalized to the average projected body area obtained from 2D images using FIJI, as previously described.^71^

### Oil Red O staining

*C. elegans* were stained with Oil Red O (ORO) as previously described.^72^ Briefly, a synchronized population of Day 1 adult animals was washed in PBST (1x phosphate-buffered saline + 0.01% Triton X-100), fixed in 40% isopropanol, then incubated with ORO working solution for 2 h at room temperature shielded from light. After washing off excess dye, animals were transferred to a glass slide, and images were taken using a Revolve microscope equipped with a color sensor (Echo).

Image analysis was conducted using ImageJ/FIJI software. ORO images were transformed into the HIB color and quantified in the (S) channel by tracing intestinal regions.

### Brood size measurement

For each condition tested, 10 synchronized L4 animals hatched on respective plates were singled onto fresh plates. Animals were allowed to lay eggs for 24 h at 20 °C and then moved to fresh plates each day until egg laying stops. Progenies laid were allowed to grow up at 20 °C and the surviving L4 larvae were counted.

### Pharyngeal pumping rate measurement

Synchronized Day 1 adult animals grown on the respective bacterial diets, with or without heme supplementation, were used for the pharyngeal pumping assay. For each condition, the number of pharyngeal contractions was counted over a 20-second interval for 15 individual animals under a stereomicroscope.

### Threshing and swim speed measurement

The thrashing assay was performed using the WormLab automated multi-worm tracking system (MBF Bioscience). In brief, Day 1 adult animals were washed off from a plate using 1 mL of M9 and transferred to an empty NGM plate. In each plate, thrashing movement of animals was recorded for 1 minute at room temperature, and the wave initiation rate and swimming speed were automatically analyzed within the WormLab software. To ensure the accurate assessment of locomotor capacity of live animals, any animals that remained stationary for more than 50% of the recording duration were excluded from the analysis. Swimming metrics are based on the previously described metrics.^73^

### MitoSOX staining

A synchronized population of Day 2 adult animals were collected to Eppendorf tubes and washed twice with M9 buffer. 5 mM MitoSOX stock solution was diluted in M9 buffer at a 1:500 ratio to make 10 μM MitoSOX working stock solution, and animals were stained for 1 h. After staining, animals were washed with M9 buffer twice, immobilized using a 5 mM levamisole solution (31742, Sigma-Aldrich) on a glass slide, and immediately imaged using a Zeiss LSM900 Airyscan super-resolution confocal microscope.

For quantification, the mean fluorescence intensity of the MitoSOX signal (red channel) was measured using ImageJ/FIJI software.

### H₂O₂ measurement using a HyPer reporter

To assess endogenous hydrogen peroxide (H₂O₂) levels in *C. elegans* upon *E. faecalis* exposure, Hyper reporter (*rpl-17p:*:HyPer) was used as previously described.^74^ In brief, Hyper is a YFP-based ratiometric H₂O₂ biosensor that undergoes a conformational change and shift its fluorescence excitation peak from 420 nm to 490 nm upon oxidation by H₂O₂.

Animals were lightly immobilized using a 5 mM levamisole solution on a glass slide and immediately imaged using a Zeiss LSM900 Airyscan super-resolution confocal microscope. Images were acquired using two distinct excitation wavelengths, 490 nm (oxidized state) and 420 nm (reduced state), with emission captured at 520 nm. The relative H₂O₂ concentration was determined by calculating the ratio of fluorescence intensity (490 nm / 420 nm) using FIJI software.

### Amplex Red Assay for measuring H_2_O_2_ generation

The levels of H₂O₂ secreted by *E. faecalis* were quantified using the Amplex Red Hydrogen Peroxide/Peroxidase Assay Kit (A22188, Invitrogen) according to the manufacturer’s instructions. Briefly, *E. faecalis* OG1RF mutants were obtained from the transposon insertion mutant library^75^ and cultured as described above. A total of 1×10^7^ CFU of *E. faecalis* and 100 μL of Amplex Red reaction buffer (50 μM Amplex Red, 0.1U/mL HRP) were added to each well of a clear flat-bottom 96-well plate and incubated for 20 minutes at room temperature. The absorbance was measured at 560 nm using Tecan Infinite M1000 plate reader (Tecan).

Absolute H₂O₂ concentrations were calculated by comparing the absorbance values to a standard curve generated with known concentrations of H₂O₂.

### *C. elegans* lifespan measurements

Lifespan assessments with live *E. faecalis* were carried out on BHI agar plates as previously described.^69^ In brief, animals were synchronized through bleaching, grown on NGM plates seeded with *E. coli* HT115 until the L4 stage, and transferred to BHI plates seeded with 10 µL of *E. faecalis,* 10 µL of *E. coli* HT115, and 80 µL of LB liquid media. For lifespan assessments with RNAi bacteria, 1 mM IPTG was added to both NGM plates and BHI plates.

For lifespan assay with dead *E. faecalis*, PFA-killed bacteria were used. The OD600 of PFA-killed E. faecalis and PFA-killed *E. coli* HT115 cultures was measured, and the cultures were mixed at a 4:1 ratio (*E. faecalis*:*E.coli*). Animals were bleach-synchronized, grown on NGM plates seeded with PFA-killed *E. coli* HT115 until the L4 stage, and transferred to NGM plates seeded with the PFA-killed *E. faecalis* and *E. coli* HT115 mixture. The bacterial plates were spotted with 100 µL of 10 mg/mL FUDR (50-91-9, Spectrum Chemical).

For cytochrome *c* supplementation during lifespan assays, 5 μM Cyt *c* was overlayed on top of the bacterial lawn, and fresh Cyt *c* was given to animals every 4 days by either transferring them to a new plate or overlaying the drug.

A total of 120 animals were monitored for mortality at intervals of 1-2 days until all of them had been assessed. Animals that crawled onto the side of the plate and died were censored and excluded from the statistical analysis.

## QUANTIFICATION AND STATISTICAL ANALYSIS

Statistical analysis was performed using the Prism 10 software (GraphPad). Data are represented as mean ± SEM unless otherwise noted. Unpaired two-tailed t test assuming Gaussian distribution was used for comparisons of two normally distributed datasets with equal variances. For two datasets with unequal variances, unpaired two-tailed t test with Welch’s correction was performed. Mann-Whitney U test was used for comparisons of two non-parametric datasets.

For multiple comparisons of normally distributed datasets with one or two independent variables, statistical significance was determined using one-way or two-way ANOVA followed by Dunnett’s multiple comparisons test (to compare the mean of multiple experimental groups against a single control) or Tukey’s test (to compare the mean of each sample with the mean of every other sample). For multiple datasets with unequal variances, Brown-Forsythe and Welch ANOVA tests followed by Dunnett’s T3 multiple comparisons test were performed. Kruskal-Wallis tests with Dunn’s multiple comparisons test were used for multiple comparisons of non-parametric datasets. Log-rank (Mantel-Cox) test was used to compare lifespans of two groups of worms. P values were used to quantify the statistical significance of the tests. For all panels, **p < 0.01; ***p < 0.001; ****p < 0.0001; ns, not significant.

The sample size per condition is indicated as “*n*=” in the figure legends. Each experiment was repeated at least three times unless otherwise stated.

## ADDITIONAL RESOURCES

### Graphics

BioRender and Microsoft PowerPoint were used for generating cartoon graphics. Prism 10 was used for generating data graphics.

---

**Figure S1.**
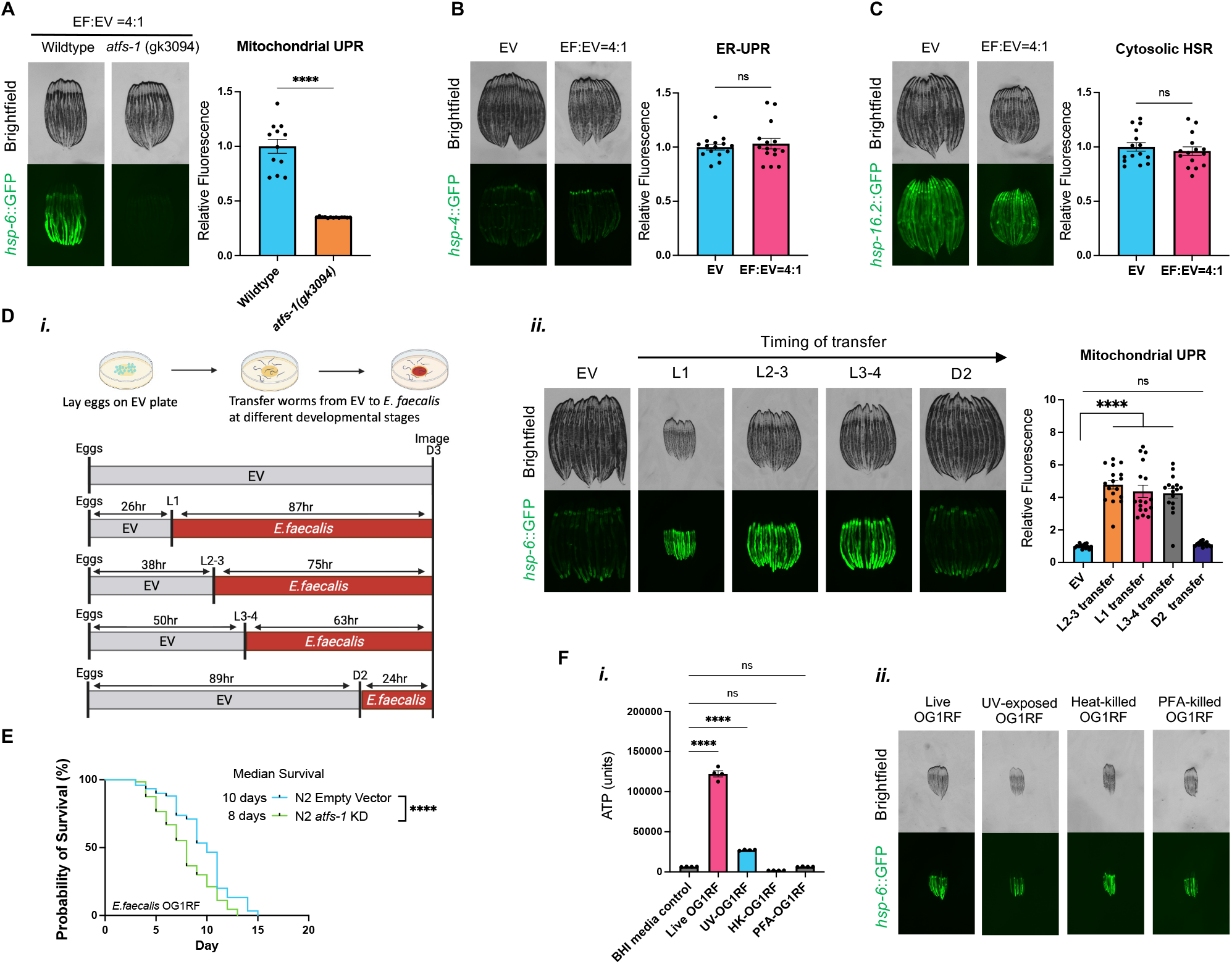
*E. faecalis* infection induces UPR^MT^ in an *atfs-1-*dependent manner, related to Figure 1. (A) Fluorescence images (left) and quantification (right) of *hsp-6p*::GFP in wildtype or *atfs-1(gk3094)* loss-of-function mutant animals hatched on a 4:1 mixed bacterial plate (*E. faecalis*:EV). *N*=2 biological replicates. (B/C) Fluorescence images (left) and quantification (right) of *hsp-4p*::GFP (B) or *hsp-16.2p*::GFP (C) in D2 adult animals hatched and grown on *E. coli* HT115 or on a 4:1 mixed bacterial plate (*E. faecalis*:EV). (D) Schematic of transfer assay (i) and fluorescence images and quantification of *hsp-6p*::GFP in D3 adult animals transferred to *E. faecalis* at the indicated timing (ii). (E) Animals were hatched on *atfs-1* RNAi or EV bacteria and grown to the L4 stage, which were then transferred to *E. faecalis* OG1RF mixed with 10% RNAi bacteria for lifespan measurement. *n*=120. (F) ATP assay (i) and fluorescence images (ii) of *hsp-6p*::GFP in D2 adult animals hatched and grown on either live *E. faecalis* OG1RF or differently inactivated *E. faecalis* OG1RF. Data are represented as mean ± SEM (A/B/C/D/F). Kruskal-Wallis test with Dunn’s multiple comparisons test (A/D); Mann-Whitney U test (B); or Student’s t test (C); Log-rank test (E); One-way ANOVA with Dunnett’s multiple comparisons test (F).

**Figure S2.**
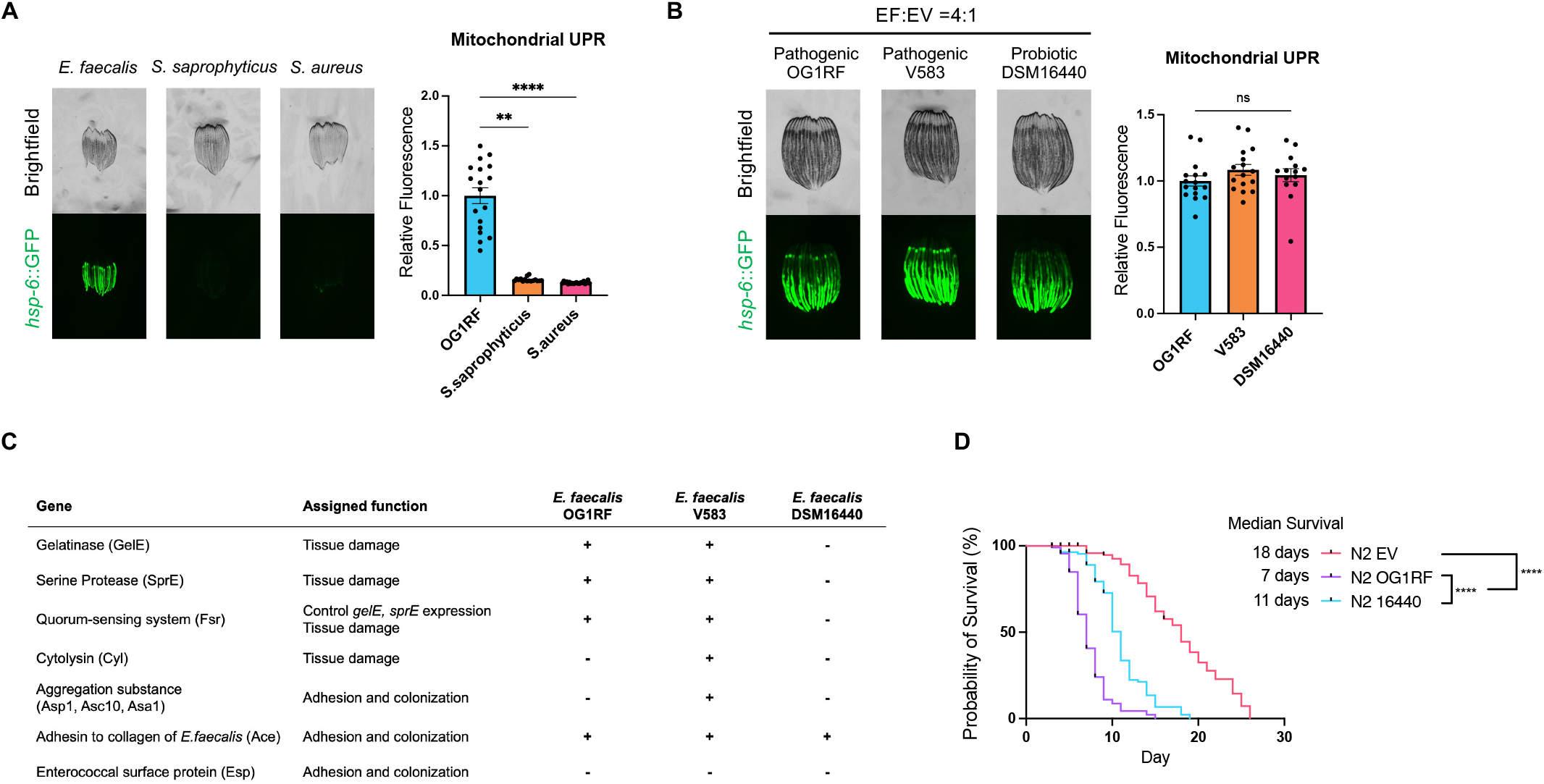
Neither virulence factors nor the growth arrest is the cause of UPR^MT^ activation in *E. faecalis*-exposed *C. elegans*, related to Figure 1. (A) Fluorescence images (left) and quantification (right) of *hsp-6p*::GFP in D2 adult animals hatched and grown on *E. faecalis* OG1RF, *S. saprophyticus*, or *S. aureus*. (B) Fluorescence images (left) and quantification (right) of *hsp-6p*::GFP in D2 adult animals hatched and grown on EV bacteria mixed with *E. faecalis* OG1RF, V583, or DSM16440 at a 4:1 ratio (*E. faecalis*:EV).. (C) Virulence factor profiles of pathogenic and probiotic *E. faecalis* strains. (D) Animals were hatched on EV bacteria and grown to the L4 stage, which were then transferred to *E. faecalis* mixed with 10% EV bacteria for lifespan measurement. *n*=120. Data are represented as mean ± SEM (A/B). Kruskal-Wallis test with Dunn’s multiple comparisons test (A); One-way ANOVA with Dunnett’s multiple comparisons test (B); Log-rank test (D).

**Figure S3.**
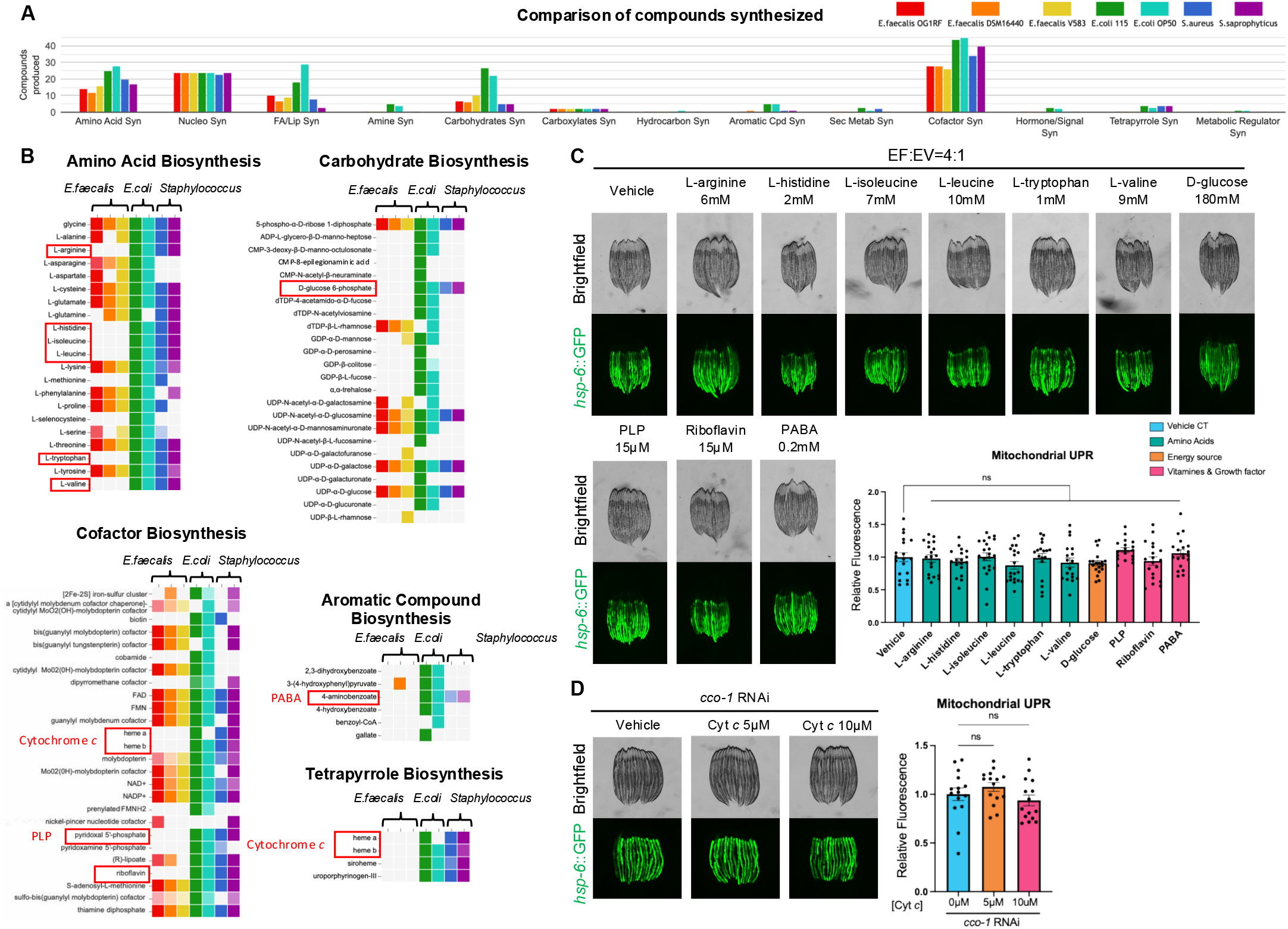
Additional details of nutrient screening, related to. Figure 2. (A) Comparative analysis of bacterial compounds synthesized by indicated bacterial strains was performed using the BioCyc database. (B) Detailed breakdown of biosynthesis pathways for amino acids, carbohydrates, cofactors, aromatic compounds, and tetrapyrroles. Bacterial compounds uniquely absent in *E. faecalis* strains are highlighted by red boxes. (C) Fluorescence images (top) and quantification (bottom) of *hsp-6p*::GFP in D2 adult animals hatched and grown on a 4:1 mixed bacterial plate (*E. faecalis*:EV) supplemented with individual CeMM component. (D) Fluorescence images (left) and quantification (right) of *hsp-6p*::GFP in D2 adult animals hatched and grown on *cco-1* RNAi bacteria supplemented with Cyt *c* 5μM or 10μM. All data are represented as mean ± SEM. One-way ANOVA with post hoc Dunnett’s test (C/D).

**Figure S4.**
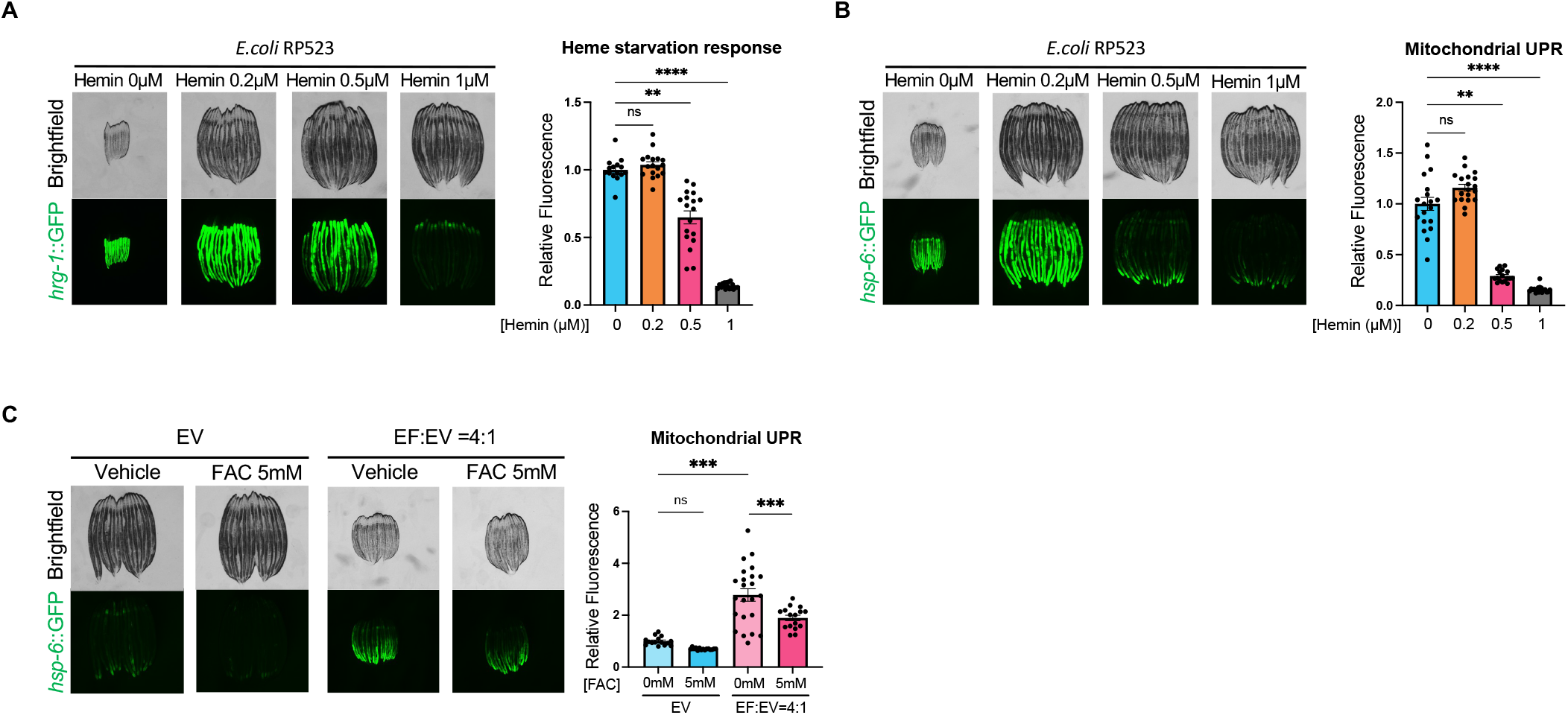
Additional evidence supporting UPR^MT^ activation caused by heme deficiency, related to. Figure 3. (A) Fluorescence images (left) and quantification (right) of *hrg-1p*::GFP in D1 adult animals hatched and grown on *E. coli* RP523 supplemented with 0, 0.2, 0.5, or 1 μM of hemin chloride solution. (B) Fluorescence images (left) and quantification (right) of *hsp-6p*::GFP in D1 adult animals hatched and grown on *E. coli* RP523 supplemented with 0, 0.2, 0.5, or 1 μM of hemin chloride solution. (C) Fluorescence images (left) and quantification (right) of *hsp-6p*::GFP in D1 adult animals grown on EV bacteria or a 4:1 mixed bacterial plate (*E. faecalis*:EV) supplemented with ferric ammonium citrate (FAC). All data are represented as mean ± SEM. Kruskal-Wallis test with Dunn’s multiple comparisons test (A/B); Two-way ANOVA with Tukey’s multiple comparisons test (C).

**Figure S5.**
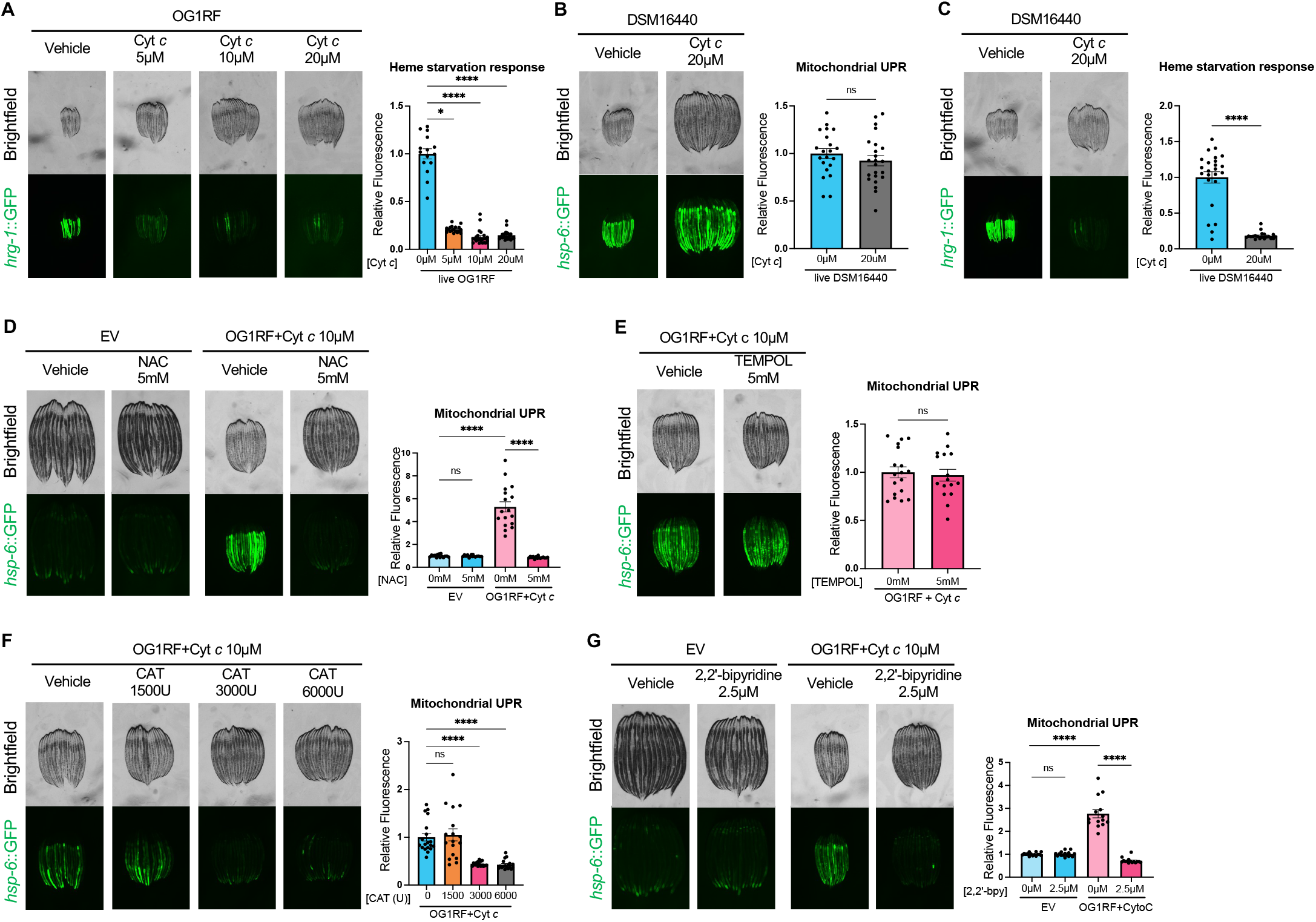
Additional evidence showing that live *E. faecalis* infection activate UPR^MT^ by inducing oxidative stress, related to Figure 6. (A) Fluorescence images (left) and quantification (right) of *hrg-1p*::GFP in D3 adult animals grown on live *E. faecalis* OG1RF supplemented with cytochrome *c* at the indicated concentration. (B/C) Fluorescence images (left) and quantification (right) of *hsp-6p*::GFP (B) or *hrg-1p*::GFP (C) in D3 adult animals grown on live *E. faecalis* DSM16440 supplemented with 20 μM cytochrome *c*. (D/E/G) Fluorescence images (left) and quantification (right) of *hsp-6p*::GFP in D3 adult animals grown on live *E. faecalis* OG1RF co-supplemented with cytochrome *c* and 5 mM NAC (D), 5 mM TEMPOL (E), or 2.5 μM 2,2’-Bipyridine (G). (F) Fluorescence images (left) and quantification (right) of *hsp-6p*::GFP in D2 adult animals grown on live *E. faecalis* OG1RF co-supplemented with cytochrome *c* and catalase at the indicated amount. All data are represented as mean ± SEM. Kruskal-Wallis test with post hoc Dunn’s test (A/F); Student’s t test (B/E); Mann-Whitney U test (C); Two-way ANOVA with Tukey’s multiple comparisons test (D/G).

**Figure S6.**
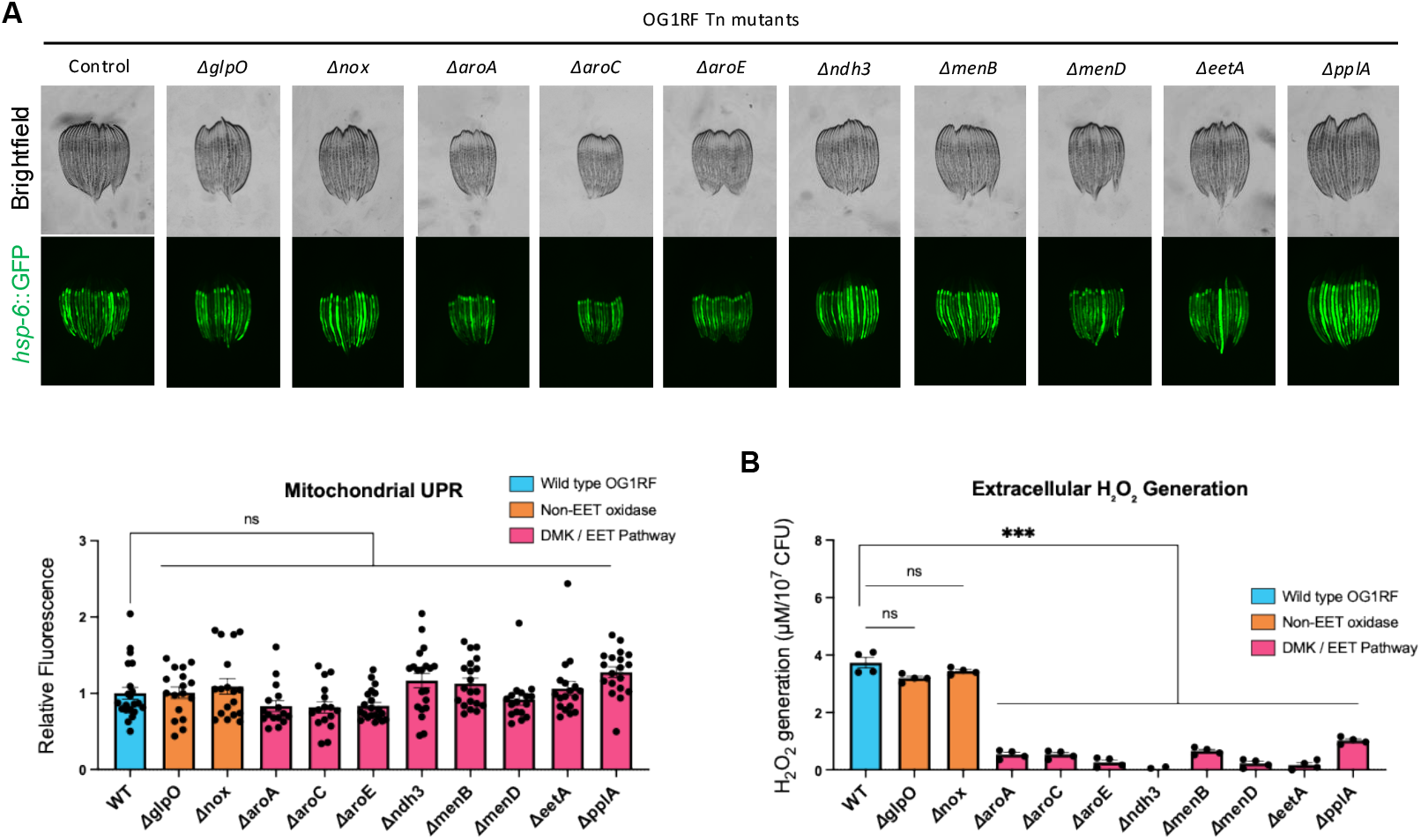
*E. faecalis*-derived H_2_O_2_ is not the cause of UPR^MT^ activation observed in live-*E. faecalis* infection, related to Figure 7. (A) Fluorescence images (top) and quantification (bottom) of *hsp-6p*::GFP in D3 adult animals grown on wildtype *E. faecalis* OG1RF (WT), non-EET oxidase mutants (*ΔglpO, Δnox*), or demethylmenaquinone (DMK) / extracellular electron transfer (EET) pathway mutants (*ΔaroA, ΔaroC, ΔaroE, Δndh3, ΔmenB, ΔmenD, ΔeetA, ΔpplA*). (B) Quantification of extracellular H₂O₂ generation by the indicated *E. faecalis* strains using the Amplex Red assay. All data are represented as mean ± SEM. Kruskal-Wallis test with post hoc Dunn’s test (A); Brown-Forsythe and Welch ANOVA tests with Dunnett’s T3 multiple comparisons test (B).

